# Long-term culture system for deep-sea mussels *Gigantidas childressi*

**DOI:** 10.1101/2021.11.24.469854

**Authors:** Claas Hiebenthal, Finn-Ole Gehlert, Mark Schmidt, Thorsten B. H. Reusch, Frank Melzner

## Abstract

The simulation of deep-sea conditions in laboratories is technically challenging but necessary for experiments that aim at a deeper understanding of physiological mechanisms or host-symbiont interactions of deep-sea organisms. In a proof-of-concept study, we designed a recirculating system for long-term culture (>2 years) of deep-sea mussels *Gigantidas childressi* (previously *Bathymodiolus childressi*). Mussels were automatically (and safely) supplied with a maximum stable level of ~60 µM methane in seawater using a novel methane-air mixing system. Experimental animals also received daily doses of live microalgae. Condition indices of cultured *G. childressi* remained high over years, and low shell thickness growth could be detected, which is indicative of positive energy budgets. Using stable isotope data, we demonstrate that *G. childressi* in our culture system gained energy, both, from digestion of methane oxidizing endosymbionts and from digesting particulate food (microalgae). Limitations of the system, as well as opportunities for future experimental approaches involving deep-sea mussels are discussed.

## 1. Introduction

Most pelagic and benthic deep-sea food webs depend on limited amounts of organic material that originates in the euphotic zone and sinks to the deeper ocean (Billett et al. 1983; Gage 2003). As a result, the biomass of most deep-sea species assemblages is low (Sanders & Hessler 1969). However, chemosynthetic deep-sea habitats such as ‘hot vents’ or ‘cold seeps’ can host a set of specialized organisms at the seafloor thriving at high abundances with biomasses of up to 30 kg m^−2^ (e.g. Paull et al. 1984; Sibuet & Olu 1998). At ‘cold seep’ sites, high-saline and cool sea water (e.g. 120 g kg^−1^ and 6.5-7.2 °C at Brine-Pool-NR-1, Northern Gulf of Mexico) seeps from the sea floor (e.g. Smith et al. 2000; Berger & Young 2006). The seeping water is usually highly enriched in reduced compounds, such as hydrogen sulfide or methane (e.g. Bergquist et al. 2005; Tunnicliffe et al. 2003) that serve as main energy sources for the organisms dwelling at such chemosynthetic ecosystems. Thus, these systems do not rely on solar energy for carbon fixation (Tunnicliffe et al. 2003). Thiotrophic, methylotrophic or methanotrophic microorganisms use chemical energy via oxidation and fixation of the reduced inorganic molecules from the seep water (McCollom & Shock 1997; Duperron et al. 2007). Some of these form bacterial mats on the sea floor which grazers like shrimp or snails feed upon (Dattagupta et al. 2007). Other chemolithoautotrophic bacteria developed close symbiotic relationships with metazoans such as polychaete worms, sponges or bivalve molluscs (MacDonald et al. 1990; Tunnicliffe et al. 2003; Cordes et al. 2009; Nishijima et al. 2010). The latter allow their animal hosts (e.g. *Riftia pachyptila*, *Bathymodiolus* spp.) to indirectly obtain energy from inorganic molecules dissolved in the seawater, either by translocation of fixed carbon products to the host (‘milking type’) or by digestion of the bacteria by their hosts (‘farming-type’ carbon transfer; Streams et al. 1997).

The deep-sea mytilid mussel *Gigantidas childressi* (subfamily Bathymodiolinae, originally described as *Bathymodiolus childressi* (Childress et al. 1986; Gustafson et al. 1998)) occurs in high abundances at cold seeps from about 550 to more than 2200 m depth on the upper as well as the lower continental shelf in the Gulf of Mexico (Nix et al. 1995; Gustafson et al. 1998; Faure et al. 2015). The mussel holobiont utilizes methane as its only energy and carbon source (Cary et al. 1988). Metanotroph γ-proteobacteria that utilize a ribulose-monophosphate pathway (RuMP) to fix carbon (‘type-1 methanotrophs’) are concentrated in vacuoles of symbiont-containing bacteriocytes in the mussel outer (water facing) gill epithelium (Duperron et al. 2007; Fiala-Médioni et al 1986; Wentrup et al. 2014). The mussels generate a ciliary-driven water current to lead methane-rich water past their gills, where methane can diffuse into the gill tissue to supply the demands of the symbionts. The methanotroph bacteria proliferate and are eventually digested by their hosts (‘farming-type’ carbon transfer, Fisher & Childress 1992; Streams et al. 1997). However, bathymodiolinid mussels do not always solely depend on carbon fixation of bacterial symbionts (Riou et al. 2010). While the host’s gastrointestinal tract is often reduced in other symbiotic holobionts, this is not the case in *G. childressi*. Here, the mussels can still filter and digest particulate organic matter (POM) from the water column (Cary et al. 1988; Page et al. 1990). At low methane availability, bathymodiolinid mussels may even obtain the majority of the assimilated carbon from POM (Martins et al. 2008; Riou et al. 2010).

Experimental work with living deep-sea animals is extremely challenging. As it is very time-consuming and costly to reach deep sea mussel reefs via remotely operated vehicles, *in situ* manipulations – such as transplantation experiments - are rare (Nix et al. 1995; Ravaux et al. 2019). On the other hand, many deep-sea animals instantly die when taken to the surface. Furthermore, it is technically challenging to recreate deep-sea pressure and water chemistry in laboratories, which makes *in vitro* experiments very difficult to realize as well (Huang et al. 2019). Such laboratory experiments, however, are crucial to understand for example energy metabolism at varying seawater methane concentrations, the relative importance of energy uptake via autotroph symbionts vs. filtration of POM, as well as the physiological mechanisms underlying host-symbiont energy transfer. Ultimately, experimental culturing systems that can simulate realistic seep environments could enable us to also study reproductive processes and larval development, behavior and chemotaxis - important aspects of bathymodiolinid biology that will allow us to better constrain larval drift models, population connectivity and, ultimately, guide deep-sea conservation efforts (Breusing et al. 2016).

Here, we describe a recirculating culturing system that allowed us to cultivate deep-sea mussels *G. childressi* for multiple years while supplying them with constant methane levels as well as POM (microalgae). We demonstrate that cultured mussels maintain and increase the ability to consume methane and are able to invest surplus energy into shell thickness growth. Tissue C and N stable isotope data reflects a mixed diet of autotrophic symbionts and POM. We discuss limitations of the culturing system as well as implications for future culture systems for deep-sea organisms.

## 2. Gigantidas childressi culture

### Mussels and sampling sites

We cultured two batches of *Gigantidas childressi* specimen for altogether 39 and 31 months, respectively. The deep-sea mussels of the first batch (208 specimen, ‘batch 1’) were collected by E/V ‘Nautilus’ using ROV ‘Hercules’ south-west of New Orleans in the Gulf of Mexico at station GC233 (‘Brine Pool NR-1’; 27° 43.4076’ N, 91°14.7798’ W) in July 13th 2014 in a depth of 650 m. At this site, cold and high-saline water seeps from the ocean floor and - due to the higher density - fills a shallow ‘brine pool’ which is surrounded by a *G. childressi* mussel bed (MacDonald et al. 1990). The mussels arrived at GEOMAR (Kiel, Germany) 14 days after being collected. The second batch of *G. childressi* (35 specimen, ‘batch 2’) was collected by MSV ‘Ocean Intervention II’ using ROV ‘Global Explorer’ north-east of the first site at station GC249 (27.72879° N, 90.51690° W) in 811 m depth on June 18th 2017.

The mussels were transported within 4 days by ship and truck to the Fisher deep sea lab at Pennsylvania State University, State College, PA. Here, the experimental animals were cultured in a recirculating seawater aquarium system (T = 8 °C, S = 34) that continuously received methane-saturated seawater. Seawater methane concentrations were not quantified during this time period. Subsequently, after 2 months, animals were shipped by plane in cooled jars (T = 5 – 8 °C, S = 34) to GEOMAR (travel time < 24h). There was no mortality during the transport of both batches.

### Description of the culture system

The long-term deep-sea mussel culture system at GEOMAR that enabled continuous culture from July 2014 to December 2019 is located in a constant temperature room and is based on a classical recirculating aquarium system (fig. 1). The system consists of three 25 L glass culture tanks that receive water via gravity feed from a 25 L header tank at a constant flow rate of 150 mL min^−1^. Water leaving the culture tanks via overflow pipes (PVC) is collected in a 100 L filter sump, where it is processed by a nitrification filter (pond external filter type 3455, EHEIM, Deizisau, Germany, filled with 12 L of HDPE bio carriers), as well as by a protein skimmer (ACF 1000V, AquaCare GmbH & Co. KG, Gladbeck, Germany). Subsequently, the culture water is led by an UV lamp (hw UV-water sterilizer Model 500, Wiegandt GmbH, Krefeld, Germany) prior to being pumped back to the header tank. During the 5.5 years of culture, water temperature was kept constant at 7.9 ± 0.2 °C and pH (NBS scale) was 8 ± 0.1. Ammonia and nitrite concentrations were assessed bi-weekly and concentrations were always < 0.05 mg L^−1^. 30% of the system water was exchanged each week. The culture water was mixed from sand-filtered Baltic Sea water (salinity 12-20, GEOMAR central facility) from Kiel Fjord and a commercial sea salt mixture (Pro Reef, Tropic Marin AG, Hünenberg, Switzerland). Salinity was set to 34.5.

**Figure 1.**
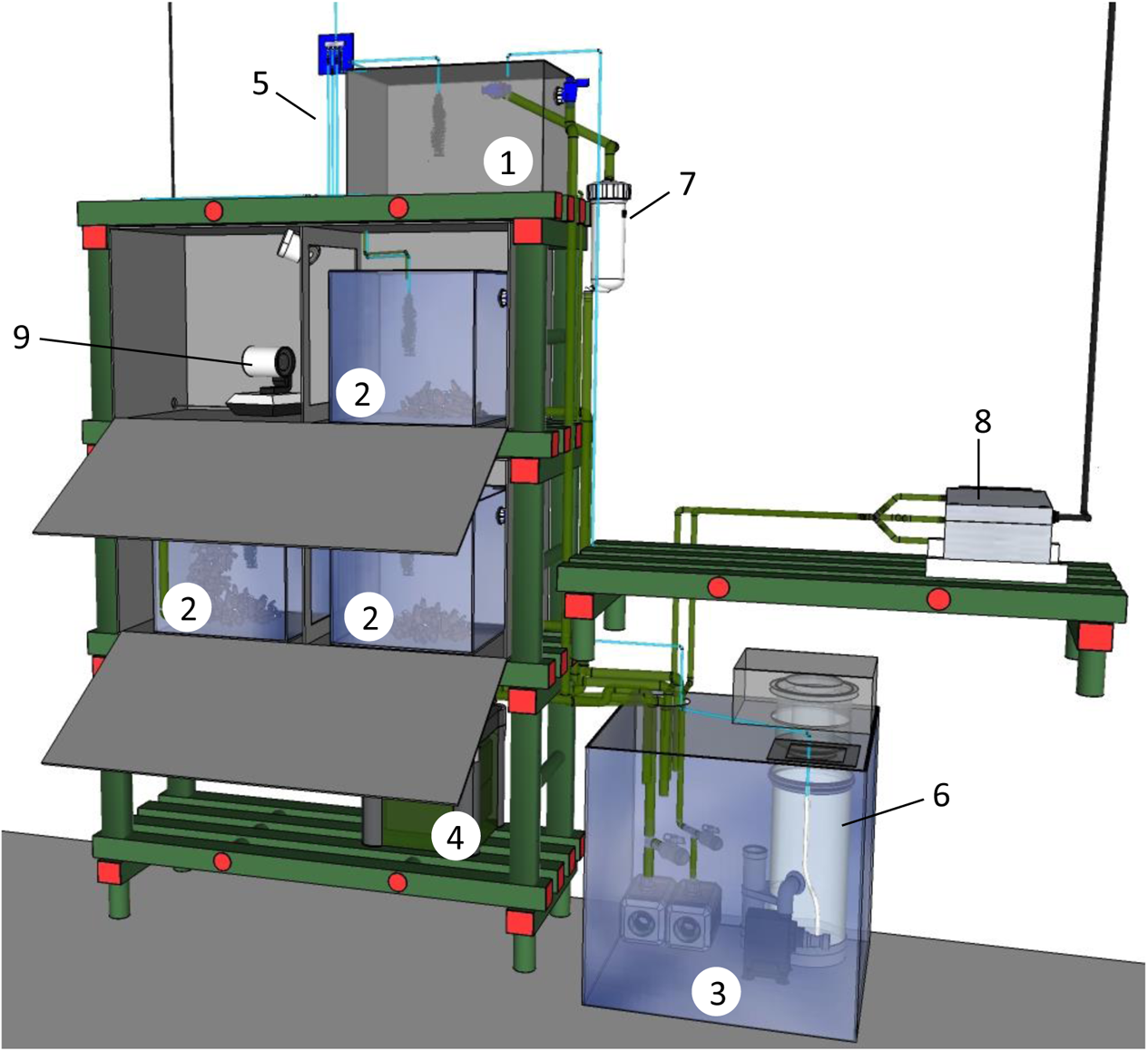
3D sketch of the 200 L water recirculating culture system to maintain deep-sea mussels *G. childressi*. Details: 1: header tank, 2: culture aquariums with deep-sea mussels, 3: filter sump, 4: biofilter, 5: gas supply line with air-methane mixture, 6: protein skimmer, 7: UV-sterilizer, 8: methane sensor, 9: IR-camera.

As *G. childressi* is sensitive to mechanical disturbances, we placed the culture tanks on rubber mats and in two PVC-boxes with openings for water and gas lines as well as flap doors at the front to access the tanks. The boxes also allowed for culturing the mussels almost full-time in complete darkness, only interrupted by short daily check-ups of the system and occasional cleaning of aquaria. An IR-camera was installed to be able to observe the mussels (fig. 1).

### Methane supply

*G. childressi* gain the majority of their energy from methanotroph bacteria in their gills. A sufficient supply with methane is therefore essential for a successful culture. At other institutes, bathymodiolinid mussels have been supplied with methane by repeated provision of methane enriched seawater e.g. by placing baskets with mussels for a limited time in a tank containing methane saturated seawater (Arellano & Young 2009, 2011; suppl. tab. 1). In a similar first feeding attempt, we initially supplied 5 L of methane-saturated seawater (corresponding to 8 mmol CH_4_) into the header tank of the 200 L culture system twice a day. This treatment, however, was labor intensive and - as methane left the system quickly – resulted in hours of methane deficiency between methane peaks (fig. 2). Under such conditions, bathymoldiolonid mussel cultures could not be maintained for more than ~6 months (Arellano & Young 2009, own trials). However, a *Bathymodiolus azoricus* recirculating culture system at LabHorta, Azores, that was continuously bubbled with methane (creating ~70 µM of methane; no continuous methane measurements) enabled mussel survival for more than 12 months (Colaço et al. 2006). Therefore, we also developed a system with a constant - and safe - methane supply to our *G. childressi* culturing system.

**Figure 2.**
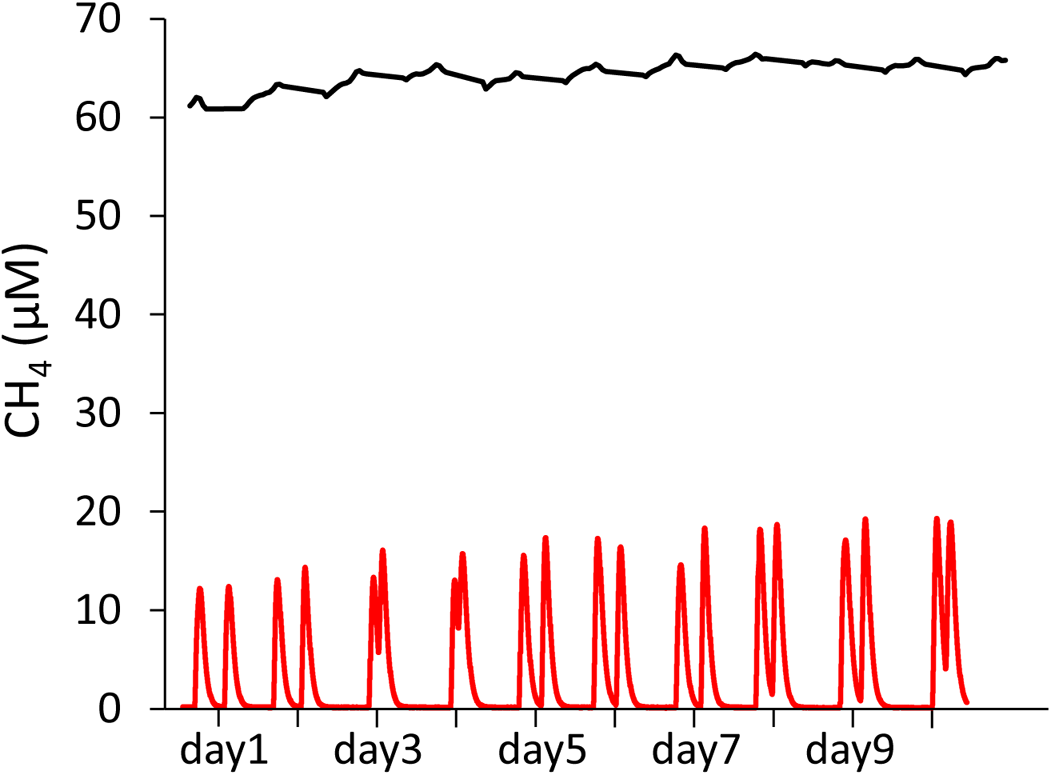
Comparison of intermittent methane feeding by repetitive addition of 5 L of methane-saturated seawater (after 15 minutes of equilibration with a gas diffusor stone, solution containing ~8 mmol of CH_4_) to the 200 L recirculation culture system (red line) with continuous methane feeding by bubbling with methane-enriched air (4% methane) from the gas mixing device (black line) as measured with a CONTROS HydroC® CH₄ sensor in the filter tank.

Central to our culture system is a methane-air gas mixing device that is placed in a room next door to the deep-sea mussel culture facility. This mixing device continuously enriches the pressurized air (‘natural’ air, continuously provided by a central supply system) led to the culture system with methane (fig. 3). The gas mixing device consists of a thermal gas flow sensor (‘red-y smart meter’, 5-500 L h^−1^, Voegtlin Instruments, Muttenz, Switzerland) and a thermal gas flow controller (‘red-y smart controller’, 0.2-20 L h^−1^, Voegtlin Instruments, Muttenz, Switzerland) that injects pure methane from a gas tank into the air flow, the amount depending on the flow rate and the desired methane level. A custom-made control unit (HTK Hamburg GmbH, Hamburg, Germany) processes the air flow rate signal from the flow sensor and defines the methane flow rate via the flow controller. Desired methane levels (0.0-4.0% methane in air) can be set using a software (EasyHTK, HTK, Hamburg, Germany) on a USB-connected PC. Gas mixture flow rate can be adjusted via a flowmeter (Q-Flow V100-140, Voegtlin Instruments, Muttenz, Switzerland).

**Figure 3.**
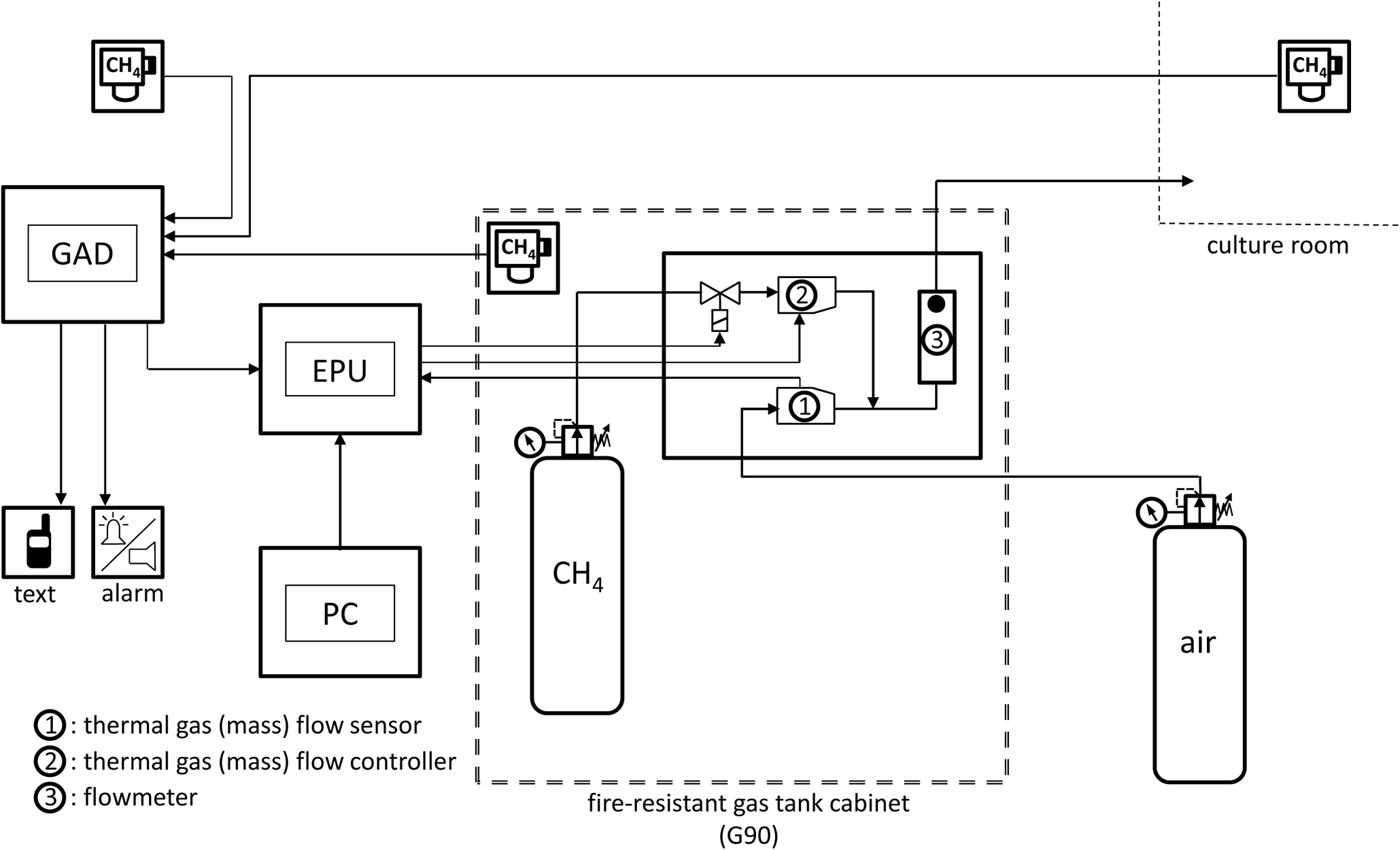
Technical sketch of the gas mixing device. The air flow is measured by the flow sensor (1) and an evaluating processor unit (EPU) controls how much methane is injected by the flow controller (2). Methane-air mixture flow can be controlled by a flowmeter (3). A gas alarm device (GAD) receives methane level signals from 3 sensors (in the fire-resistant gas tank cabinet, in the gas mixture room and in the deep-sea mussel culture room) and triggers alarm signaling, text messaging and closing of the magnetic valve (cutting-off methane flow) when alarm levels are reached.

The gas mixture is then led in a gas-tight polyurethane line (PUR C 98A) from the gas mixing device room to the *G. childressi* culture system room. Both, the header tank and the three culture tanks of the culture system are equilibrated with the gas mixture using diffusor stones at ~1.7 L gas mixture min^−1^. Also, the protein skimmer is supplied with the methane-air gas mixture instead of the methane-poor culture room air. The total gas flux of 4% methane-air mixture to the recirculation culture system is 400 L h^−1^. Methane concentrations in the culture system water are continuously monitored with a flow-through instrument that can determine seawater methane concentration with an accuracy of ± 0.1 µM (CONTROS HydroC® CH₄ FT, 4H-JENA engineering, Kiel Germany). Culture seawater is pumped to the methane sensor at a rate of 2 L min^−1^ (EHEIM universal pump 1200, EHEIM).

With the described system, we are able to maintain maximum seawater methane concentrations in the aquarium system at ~60 µM (suppl. fig. 1, fig. 5) which corresponds well to an expected methane equilibrium concentration of the injected 4% methane in air mixture of 62.14 µM (using Bunsen solubility coefficient at salinity 34 and 8 °C; Wiesenburg and Guinasso 1979). These values are higher than lower concentrations measured within *Gigantidas* reefs in the Gulf of Mexico (‘Bush Hill’: 20-56 µM CH_4_, ‘Green Canyon’: 42-10744 µM CH_4_, Nix et al. 1995, Smith et al. 2000), similar to those provided by Colaço et al. (2006) to their *Bathymodiolus azoricus* culture (~70 µM CH_4_) and allowed us to culture *G. childressi* for years.

**Figure 5.**
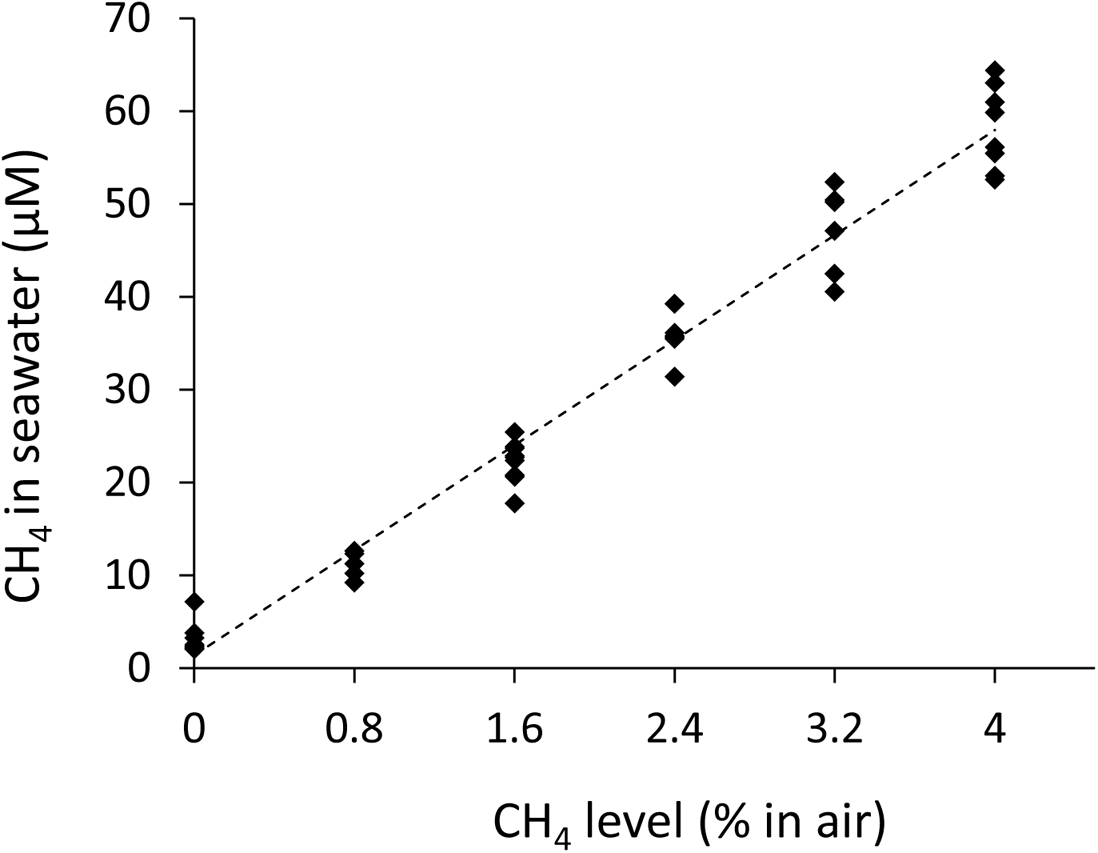
System performance during the mussel valve gape activity experiment: Daily averages of measured CH_4_ concentrations in the recirculating aquarium system at six different CH_4_ concentrations (%) in the air used to equilibrate the seawater. CH_4_ (µM) = 14.149 %CH_4_ + 1.3907; R² = 0.98, regression: F = 1294.2, p < 0.001; n = 6.

### Safety measures

As our deep-sea mussel culturing system can be run close to the lower explosion limit for methane in air, a set of safety measures was installed to exclude the formation of an explosive atmosphere in a non-explosion safe environment:

i. The maximum methane level that can be produced by the gas mixing device is (software-wise) restricted to 4%, which is below the explosion limit of 4.4% methane in air.
ii. The gas tank as well as the mixing device’s gas leading parts (flow sensor, flow controller, flowmeter) are placed inside a fire-resistant gas tank cabinet (G90, in accordance with EN 14470-2, fire resistance of 90 minutes). Only the processor unit is placed outside, and only the gas mixture with a maximum of 4% methane leaves the cabinet in gas-safe lines (no pure methane).
iii. Methane sensors are placed in the constant temperature room (*G. childressi* culture room), in the gas mixing room and in the fire-resistant gas tank cabinet containing the mixing device. A central gas alarm device raises an alarm when two alarm levels are reached: *alarm level 1* at 0.88% methane (20% of lower explosion limit, flash light) and *alarm level 2* at 1.76% methane (40% of lower explosion limit, flash light and alarm sound). At both alarm levels text messages are sent to operating personnel.
iv. At *alarm level 1* a magnetic valve between gas bottle and flow controller is closed and, hence, the methane flow is interrupted.
v. Power failure also causes the magnetic valve to close, and the alarm device to send a text message to operating personnel.
vi. Operating personnel controls methane consumption rates of the mixing device and methane concentration in the culture system water daily to detect potential leaks.

Costs for setting-up the entire deep-sea mussel culture system installed in 2013-2014 were ~17,000€ for the culturing system (including methane sensing instrument HydroC® CH₄ FT), another ~17,000€ for the gas mixing system, and 7,000€ for the gas safety surveillance system. Not included are repeated occurring costs for maintenance and methane consumption (at 400 L h^−1^ production of 4% methane-air mixture: ~10,000 L methane, corresponding to one 50 L 200 bar cylinder month^−1^).

### Feeding with micro-algae

Apart from consumption of methanotrophic symbionts, *G. childressi* can obtain energy from filtered organic particles as well (Cary et al. 1988; Page et al. 1990). Thus, our deep-sea mussels were also fed 5 days a week with 450 mL *Rhodomonas* sp. microalgae suspension (with ~2 x 10^6^ cells mL^−1^, corresponding to ~4500 cells mL^−1^ after being diluted in the 200 L-culture system). This cryptophyte is very rich in polyunsaturated fatty acids (Renaud et al. 1999) and is permanently cultured at GEOMAR as nutritious food for filter feeding organisms in artificial seawater supplied with ‘Walne’s medium’ including macro-nutrients, vitamins and trace elements (Walne 1970).

## 3. Experimental observations and growth of cultured deep-sea mussels

Two sets of experiments were conducted during the culture system development process. In the first set, *G. childressi* from batch 1 were cultured with 20-30 µM methane in seawater to obtain first insights into the mussels’ energy uptake. Based on these findings we decided to increase the seawater methane concentration and to start culturing mussels from batch 2 under three different nutritional regimes during the second set of experiments: i) exclusively methane (60-70 µM) = ‘m’, ii) methane (60-70 µM) and microalgae (*Rhodomonas* sp.) = ‘a+m’, and exclusively microalgae (*Rhodomonas* sp.) = ‘a’. To achieve the third nutrition treatment (without methane) one of the aquaria was separated from the main culturing system and the mussels were fed with *Rhodomonas* sp., only (250 mL d^−1^). The water of this aquarium was treated with a nitrification filter (ecco pro 200, EHEIM) and a protein skimmer (Power Skimmer WT350, Sander, Uetze-Eltze, Germany) placed in an extra 25 L-filter sump. The *G. childressi* in one of the aquaria of the main system continued to receive *Rhodomonas* sp. (250 mL d^−1^ poured directly into the aquarium) and methane - while phytoplankton addition was stopped in the other (‘m’)-aquarium. The 3-step filtering of seawater in the main recirculation system (by nitrification filter, protein skimmer and UV-clearer) prevented not-consumed microalgae from being transported to the ‘m’-treatment aquarium. Aquaria with mussels that were fed with microalgae received 125 mL *Rhodomonas* sp. suspension twice a day. The second set of experiments aimed at a deeper understanding of *G. childressi* energy acquisition modes (tab. 2).

## Methods

*Condition of cultured deep-sea mussels*: To measure the general condition of cultured *G. childressi*, the condition index (C_i_) was used as the quotient of ash-free dry weight and inner mussel shell volume (Smith 1985; Davenport & Chen 1987; Smith et al. 2000, eq. 1).

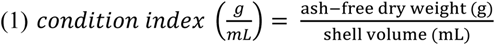

To assess the C_i_, the mussels’ soft tissue was removed from the shells, dried for 24 hours at 80 °C and the dry weight was measured. Subsequently, the tissue was incinerated at 500 °C to measure the ash weight which was subtracted from the dry weight to obtain ash-free dry weight (AFDW).

The inner shell volume was measured indirectly via the weight of fine silica sand (grading 50-1000 µm) that filled the shell: The mussel shells were filled with water to ensure a balanced position and glued to glass slides. After drying, the shells were weighed with and without sand. A calibration curve produced using 1 to 40 units of a standardized cup of exact 0.2602 mL (suppl. fig. 2) allowed us to estimate shell volume from the sand’s weight (eq. 2):

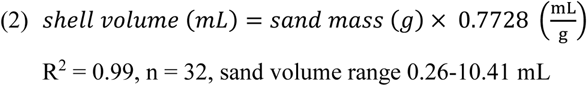

Shell lengths of the sacrificed *G. childressi* were also measured to correlate the mussels’ condition with their size. C_i_s of sub-samples of cultured deep-sea mussels were assessed three times: once with mussels of batch 1 (12 specimens in August 2016), twice with mussel of batch 2 (3 specimen in August 2017, 12 specimens in November 2019).

*Growth of cultured deep-sea mussels*: Shortly after the start of the *G. childressi* culture efforts with mussel batch 1 on September 11th 2014, the animals were stained with the fluorescent dye calcein (Sigma C0875). Calcein can be excited at 488 nm and emits at 515 nm and is rapidly incorporated into the shell of molluscs (Holcomb et al. 2013, Ramesh et al. 2017). Hence, shell length and thickness growth rates can be followed using a fluorescence stereo microscope (Nedoncelle et al. 2013). Calcein was added to the culture system (50 mg L^−1^) and subsequently - over the course of 2 months - diluted via regular water changes, allowing the mussels to incorporate the dye into newly grown shell material. In August 2016, twelve *G. childressi* individuals were sacrificed to analyze growth of calcein stained animals. Shells were analyzed using a Leica M165 FC stereo microscope at magnifications between 40–120 fold using a GFP filter set (excitation: BP 450-490 nm, emission: LP 515 nm) and EL6000 light source (Leica Camera AG, Wetzlar, Germany). Shells were first analyzed for longitudinal growth, then manually broken along the length axis and imaged in cross section at the leading edge (see fig. 4). Images were analyzed using Leica LAS X software. To follow *G. childressi* growth rates macroscopically as well, all mussel shells were furthermore marked with small scrapings at the outer shell margin (using a rotary tool to remove the periostracum at these locations without perforating the shell).

**Figure 4.**
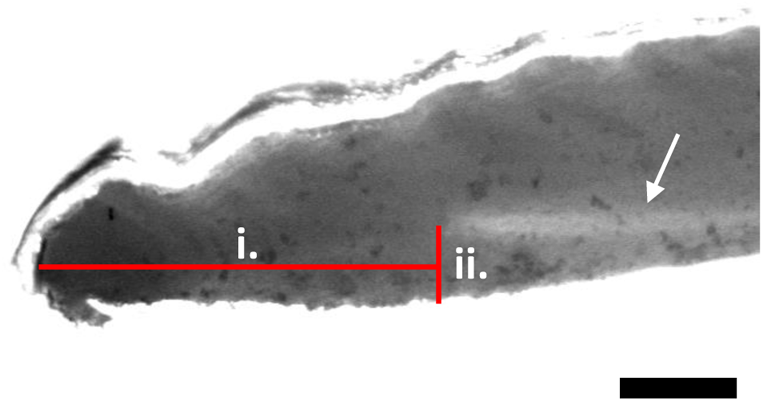
Shell length (i.) and thickness (ii.) growth measurements of cultured *G. childressi* from batch 1 after 23 months of culture. Calcein-marked growth bands (arrow) imaged with a Leica M165 FC fluorescence stereo microscope. Scale (black bars): 100 µm.

*Methane and oxygen consumption of deep-sea mussels*: To measure *G. childressi* gas consumption rates the animals were placed into gas tight closed respirometer vessels (500 mL glass bottles) that were prepared with oxygen sensor spots (SP-PSt3-NAU, PreSens Regensburg, Germany; resolution ± 0.1% at 20.9% O_2_ or ± 0.04 mg L^−1^ at 9.1 mg L^−1^) to measure oxygen concentrations with a four-channel Oxy-4 micro instrument (PreSens) via optical fibers (e.g. Vajedsamiei et al. 2021). Temporal resolution of oxygen measurements was 4 h^−1^ (suppl. fig. 7). The time animals needed to recover following transfer into respirometers (i.e. the time until they started to open their valves to filter water) varied between 1 and 3 hours and could be estimated from the point of time when respirometer oxygen concentrations started to decline. Periods of 2 or more consecutive measurements without decreasing oxygen content indicated shell closure and cessation of filtration and gas exchange rates (suppl. fig. 7; e.g. Vajedsamiei et al. 2021). Measurements were stopped after 8 hours.

Methane consumption rates were measured as two-point measurements in parallel to the respiration measurements using water samples from the respirometers. For this purpose, water samples were taken (in gas-tight 20 mL brown glass bottles) at the beginning and at the end of the measuring period. Gas (air) bubble-free water samples were poisoned immediately after sampling with 100 µL saturated mercury-chloride solution. During the measurement period the measuring chambers remained closed and without any gas (air) bubbles. Methane concentrations were measured later on a HP 5890 gas-phase chromatograph (Hewlett-Packard, Santa Clara, USA) using the headspace technique (Bange et al. 2010). During the measurements of batch 1 mussels and the first measurements of batch 2 mussels, activity of mussels could be estimated from parallel respiration measurements, see above (suppl. fig. 7). During the 2^nd^ measurements of methane consumption of batch 2 mussels, however, the mussels’ activity (open vs. closed valves) was individually monitored using cameras (QuickCam®, Logitech, Lausanne, Switzerland, connected via USB to a PC). Thus, we were able to normalize methane consumption to the amount of time animals were active (open valves, high respiration rates).

A water filled respirometer during each run was used as the control for bacterial oxygen and methane consumption during measurements and bacterial control respiration rates were always <1% of experimental animal respiration rates. Methane and oxygen consumption rates (MCH_4_ and MO_2_, respectively) per g soft body mass were calculated as the difference of gas concentrations at the start (C_start_) and at the end of the measurement (C_end_), multiplied by the volume of the measuring chamber and followed by subtraction of control (microbial) respiration rates (eq. 3):

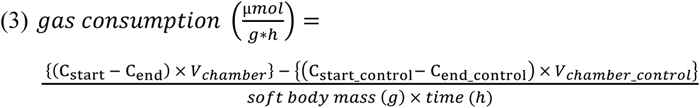

The soft tissues mass of the utilized mussels could not be assessed directly as 20-50% of the soft tissue (the gills) were sampled for symbiont density analyses. Total soft tissue could not be weighed before gill sampling as this would have corrupted the samples. Hence, soft tissue masses were indirectly estimated from a total weight – wet weight relation obtained from mussels sacrificed for C_i_ analyses (eq. 4, suppl. fig. 3):

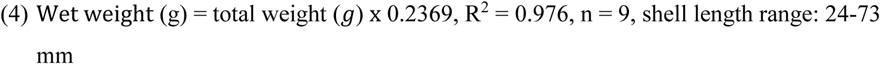

Methane and oxygen consumption measurements were conducted with both batches of animals at different time periods: animals of batch 1 following culture at 20-30 µM methane (n = 12) and animals of batch 2 following culture under the three different nutrition regimes: ‘m’, ‘a+m’ and ‘a’ (n = 3 each). Another set of measurements of the same batch 2 mussels’ methane consumption was conducted after another 19.5 months of the respective treatments to explore possible acclimation effects of *G childressi* or symbionts over time (tab. 1).

**Table 1.**
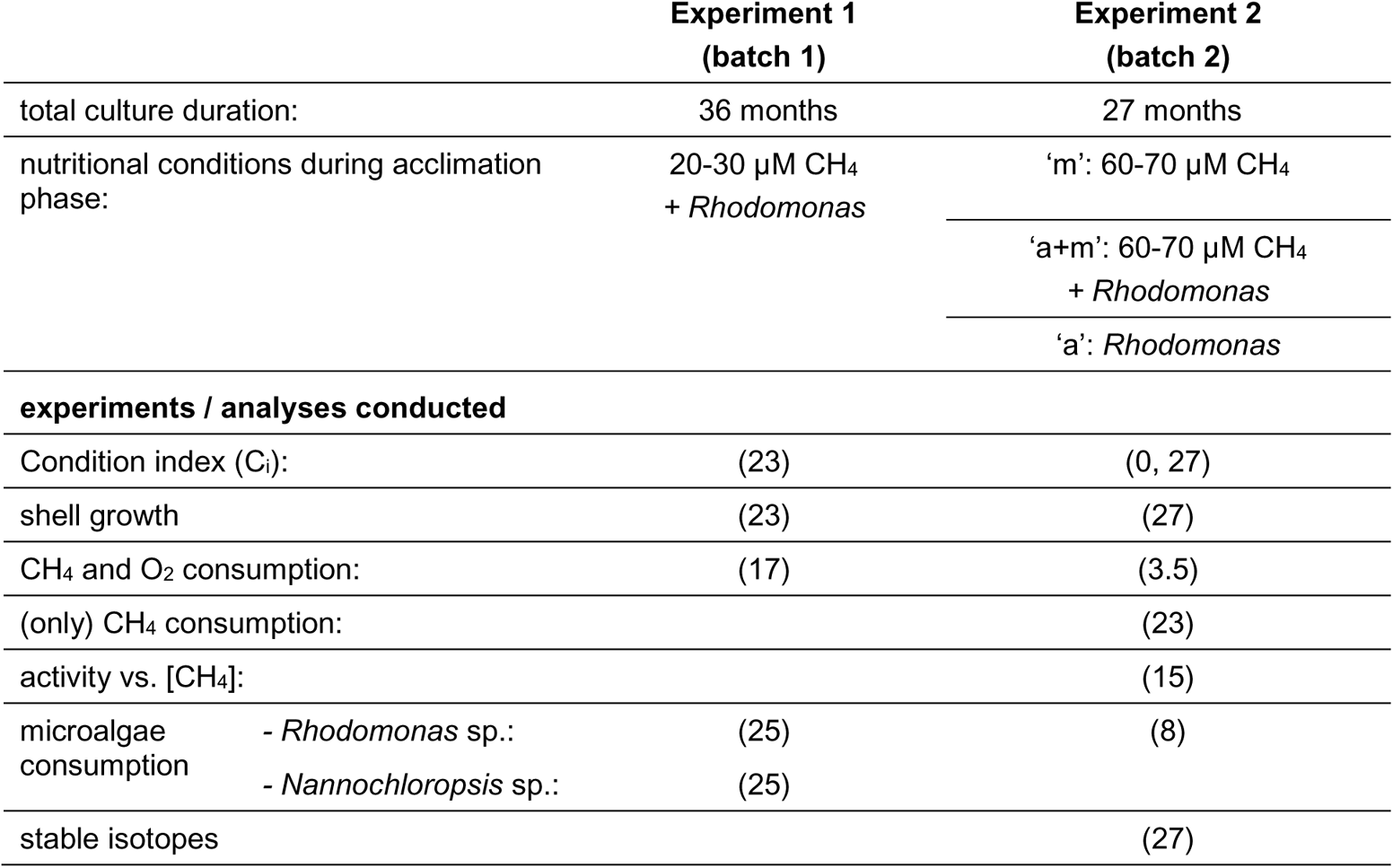
Overview of *G. childressi* experiments with mussels from batches 1 and 2. Numbers in brackets indicate the time (t, number of months) of culture after which the respective experiment was conducted. t = 0 refers to time of arrival at GEOMAR, which corresponds to ca. 0.5 and 2 months post-capture for batches 1 and 2, respectively.

*G. childressi filtering behavior in relation to water methane concentration*: Repeated IR-camera observations of cultured *G. childressi* suggested that the deep-sea mussels react to changes in culture seawater methane concentrations with increased or reduced filtering activity. To test whether these observations can be repeated under controlled conditions, a simple experiment was performed in one of the aquaria: nine batch 2 *G. childressi* individuals from the three treatments were placed into a perforated acrylic rack with multiple chambers to constrain animals (suppl fig. 4). Strong aeration and an additional circulation pump (Koralia, 900, HYDOR srl, Bassano del Grappa, Italy) increased water movement in the aquarium and into the acrylic rack. Mussel filtering behavior was observed with the IR-camera while culture system seawater methane concentrations were raised or lowered. During the experimental observations, methane concentration was raised in a stepwise manner from 0 to ~60 µM in daily increments, and then, subsequently, reduced stepwise back to 0 µM (suppl. fig. 5). This procedure was repeated three times. The IR-camera took pictures at a rate of 30 h^−1^. The pictures were analyzed after the experiment was finished by creating time lapse videos. Videos were observed and periods with opened valves were quantified for each animal. Mussels were assumed to be actively filtering with opened valves. *G. childressi* filtering activity was expressed as minutes of activity per hour.

*Particulate food consumption of deep-sea mussels*. In order to estimate the fraction of energy that can be obtained through consumption of particulate organic matter, we conducted two experiments (tab. 1) similar to those published by Page et al. (1990) using microalgae as food source.

In the first experiment, six batch 1 *G. childressi* were individually placed into 500 mL experimental units (EUs). EUs were glass bottles filled with seawater and continuously aerated with methane-air mixture (3% methane; corresponding to 40-50 µM) via diffusor stone placed in the center of each EU to aid internal mixing. Two further EUs were used as controls and did not contain animals. Before microalgae were added to the EUs, 1 mL samples were taken to determine background particle concentrations. Microalgae culture concentrations were measured using a flow cytometer (Accuri 6, BD Biosciences, San Jose, CA). Cell concentrations at the start of the experiments were set to ~7000 cells mL^−1^ (Foster-Smith 1975; Page et al. 1990). Microalgae were allowed to be evenly distributed by the circular currents in the EUs. Hence, first samples (t_0_) were taken after 15 minutes when mixing was complete. Further samples were taken after 3, 7, 11 and 24 hours. At each sampling time point, 1 mL of seawater was taken from each EU and microalgae cell concentrations were measured on the flow cytometer. The experiment was conducted first with *Rhodomonas* sp. (cell size: 5-8 µm) and subsequently with *Nannochloropsis* sp. (cell size: 2-4 µm) to test for possible feeding preferences of *G. childressi*. Clearance rates (R_C_) were calculated after Coughlan (1969; Jacobs et al. 2015):

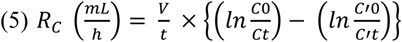

Where ‘V’ is the water volume (mL), ‘t’ is the time since start of the experiment (h), ‘C_0_’ and ‘C_t_’ are the cell concentrations of EUs with mussels filtering at t_0_ and t, and ‘C’_0_’ and ‘C’_t_’ of controls at t_0_ and t, respectively (cells mL^−1^).

In the second experiment, we used each three batch 2 *G. childressi* specimen from the three different nutritional treatments. The mussels were placed in 500 mL EUs that were equilibrated with 4% methane-air mixture (corresponding to 60-70 µM) as described above. Three additional EUs were used as controls. Cell concentrations were measured with a Coulter Counter (Z2 Coulter particle count and size analyzer, Beckman Coulter TM, Brea, CA) which allowed for quicker data acquisition. Microalgae density was set to ~7000 cells mL^−1^ at the start of the experiments. Samples were taken after 15 minutes (t_0_) and after 3, 7 and 11 hours. Microalgae consumption rate of *G. childressi* from the ‘a’ treatment group were conducted twice: Once in water equilibrated with 4% methane-air mixture and once in water without methane.

*Stable isotope (δ^15^N, δ^13^C) analyses of G. childressi cultured under different nutrition regimes*. We used batch 2 deep-sea mussel specimens acclimated to all three nutritional regimes (tab. 1) for stable isotope (δ^15^N, δ^13^C) analyses. In addition, three specimens of batch 2 were sampled directly when they arrived in our lab.

The soft tissues of the mussels were frozen at −20 °C, subsequently freeze-dried until all water had left the sample and ground to powder. Total particulate carbon and nitrogen content and isotopic compositions were analyzed according to Sharp et al. (1974). 0.5 mg of each sample was weighted into tin cups. Inorganic carbon was removed by addition of 0.2 µl of 10% HCl. After drying the samples, the procedure was repeated once more and the cups were closed and analyzed. The samples were then combusted in a CN-analyzer (Flash IRMS EA IsoLink CN Thermo Fisher) connected to a DELTA V Adv MS isotope ratio mass spectrometer (Thermo Fisher Scientific, Waltham, MA). Isotopic rations δ^15^N and δ^13^C were calculated as:

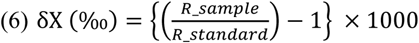

Where ‘X’ is ^15^N or ^13^C and ‘R’ is ^15^N/^14^N or ^13^C/^12^C, respectively. As primary standards, pure N_2_ and CO_2_ gases were used, calibrated against IAEA reference standards N1, N2, N3, NBS22, and USGS24. In addition to the standard calibration at the beginning of each run, standard materials (caffeine, peptone, and acetanilide) were also included within runs to identify any drift and ensure accuracy and full combustion of the samples during analysis. Overall analytical precision was ± 0.1‰ for δ^15^N and δ^13^C.

*Rhodomonas* sp. culture samples were gently centrifuged for 15 minutes at 800 rpm. and the supernatant was discarded. The concentrated cell suspension was frozen at −20 °C, subsequently freeze-dried and ground to powder as described above.

To determine stable carbon isotope fractionation between methane and cultured deep-sea mussels, two samples of the 4% methane-air mixture were taken with Tedlar® gas sampling bags. Methane was separated from other gases by gas chromatography and subsequently converted to CO_2_ in a combustion oven, which is connected to a Thermo MAT 253 continuous-flow isotope ratio mass spectrometer (CF-IRMS, Thermo Fisher Scientific, Waltham, MA). The measured ^13^C/^12^C-ratios are given in the common delta notation versus Vienna PeeDee Belemnite (VPDB). Reproducibility of ±0.3‰ (2 SD) is based on repeated measurements of laboratory reference gas.

Finally, shifts of soft tissue stable isotope composition after transplantation to a site with differently fractionated N or C can be interpreted as turnover as a measure of metabolic rates (Dattagupta et al. 2004). In that sense, annual carbon turnover rates of whole *G. childressi* soft tissue were calculated as:

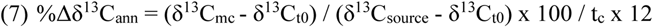

where ‘m_c_’ are cultured mussels, t_0_ are the mussels sampled right after arrival at GEOMAR and ‘t_c_’ represents the time (duration) of the culture [months].

### Results

#### Performance of the methane supply system

The continuous air-methane mixture supply to our recirculating aquarium system with up to 4% methane allowed for extended near-equilibration CH_4_ concentrations above 60 µM and, hence, a long-term culture of *G. childressi* (figs. 2 and 4, suppl. fig. 1). Deviations from optimal (stable) CH_4_ concentrations were caused intentionally due to major improvements of the gas mixture supply (improvements of gas flow volume in system in February 2015 and July 2017) and maximal %CH_4_ level (July 2017) could be realized, or when experiments were conducted that had to take place in the same culture system (suppl. fig. 1). During such experiments, desired CH_4_ levels could be selected by changing the %CH_4_ in the air-methane mixture bubbled into the system between 0% and 4% CH_4_ (fig. 5). More seldom (6 time over the entire culture period) technical malfunctions caused further reductions of methane concentrations. Overall, these intentional and non-intentional incidents together accounted for a significant amount of variation in the seawater methane concentration, most prominently so during the phase October 2017 – March 2019, where the methane concentration was 57.3 µM ± 19.7 SD (suppl. fig. 1). However, when excluding the phases of experimental work during that phase, the methane concentration was 65.7 µM ± 5.4 SD demonstrating that the system is capable to maintain very stable seawater methane concentrations.

#### Condition and Growth

Batch 2 *G. childressi* acclimated to CH_4_ (‘m’) or CH_4_ and microalgae addition (‘a+m’), maintained condition indices similar to those of the freshly imported animals (‘t_0_’) even after more than two years of culture (fig. 6a). Only smaller *G. childressi* specimen of batch 1 that were cultured at less optimal CH_4_ concentrations and batch 2 mussels that were exclusively fed with microalgae (‘a’, no CH_4_) were characterized by lower condition indices. In the latter treatment, only half as many *G. childressi* survived when compared to treatments with CH_4_ supply (fig. 6b). Data from animals freshly (<1 day post capture) collected at the same site as our batch 1 mussels (GC 233: ‘Brine Pool NR-1) indicate ca. 2-fold higher condition factors than can be reached in the field (Smith et al. 2000).

**Figure 6.**
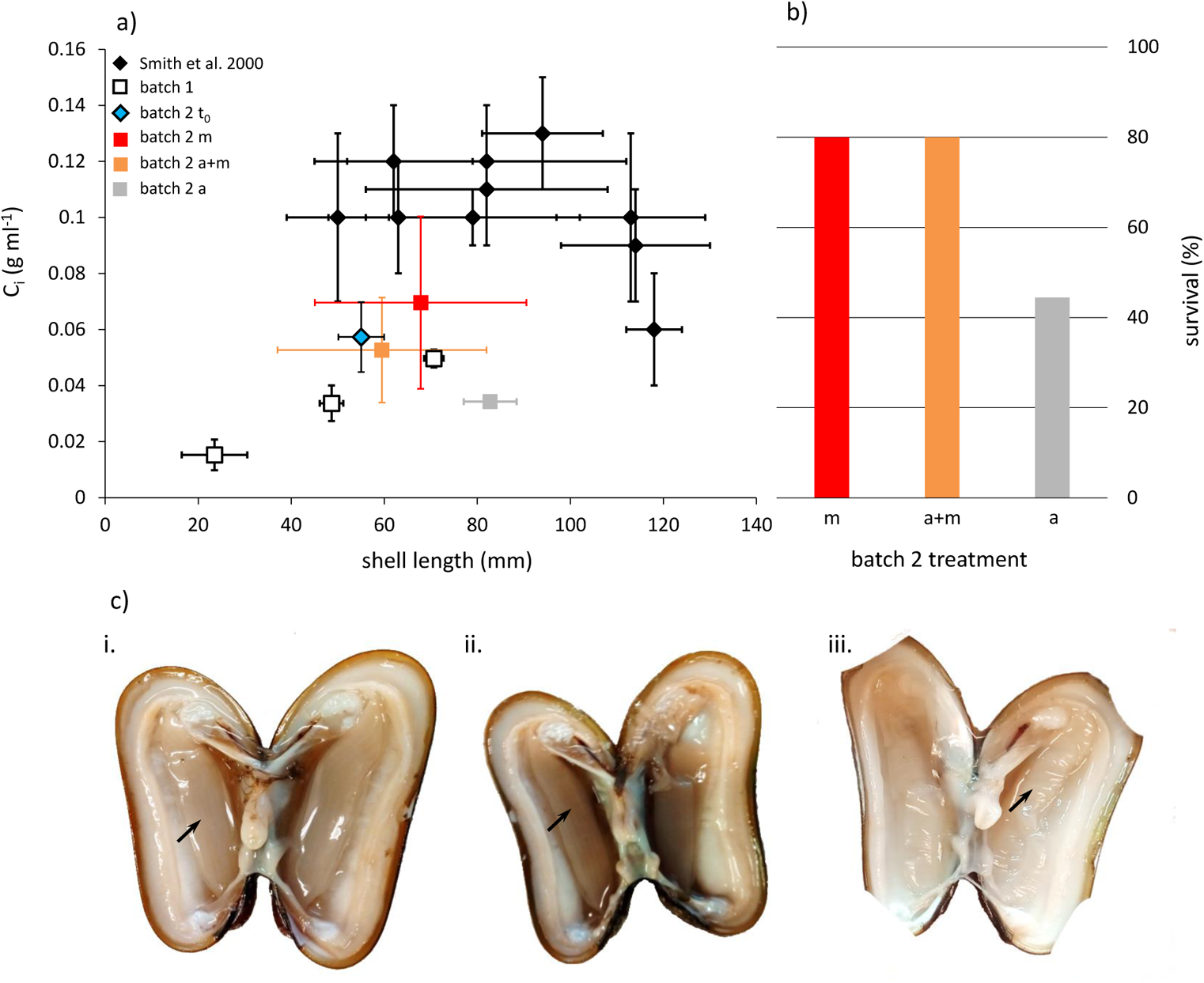
Condition (a, c) and survival (b) of batch 1 and 2 *G. childressi*. (a) Condition of ‘t_0_’ mussels was determined when batch 2 mussels arrived in Kiel. *G. childressi* from batch 1 (cultured from July 2014 to August 2016) as well as from three different nutritional treatments of batch 2 (cultured from August 2017 to November 2019, ‘m’: CH_4_, ‘a+m’: algae and CH_4_, ‘a’: algae) were sampled after 25 and 27 months of culture, respectively. Data in a) are mean ± SD. (b) Mortality of animals in treatments ‘m’, ‘a+m’ and ‘a’, that were not sacrificed for other analyses (initial animal numbers: 9-10). (c) Examples of freshly sacrificed batch 2 mussels from different nutritional treatments (CH_4_: ‘m’, algae + CH_4_: ‘a+m’, only algae: ‘a’) illustrate white - almost transparent - gills (arrows) of mussels fed without methane.

Good condition indices in batch 2 animals that received methane as energy source (‘m’ and ‘a+m’) suggest that energy supply was sufficient to prevent consumption of body mass during the culturing experiment. The observation of a net shell thickness increase in all examined specimen furthermore indicates that surplus energy could be invested into anabolic processes (fig. 7). Shell thickness increased exponentially with mussel size (fig. 7b). On the other hand, shell length increase remained low: calcein markings of 10 individual batch 1 shells revealed an increment of shell lengths between 0 to 1700 µm during the culturing period of 21 months (fig. 7a), independently of mussel size. No shell length increments were observed in two animals. Only one mussel grew 1.7 mm in shell length, all other animals grew by 400µm or less. No shell length growth could be detected in batch 2 animals, which were marked via shell (periostracum) scrapings.

**Figure 7.**
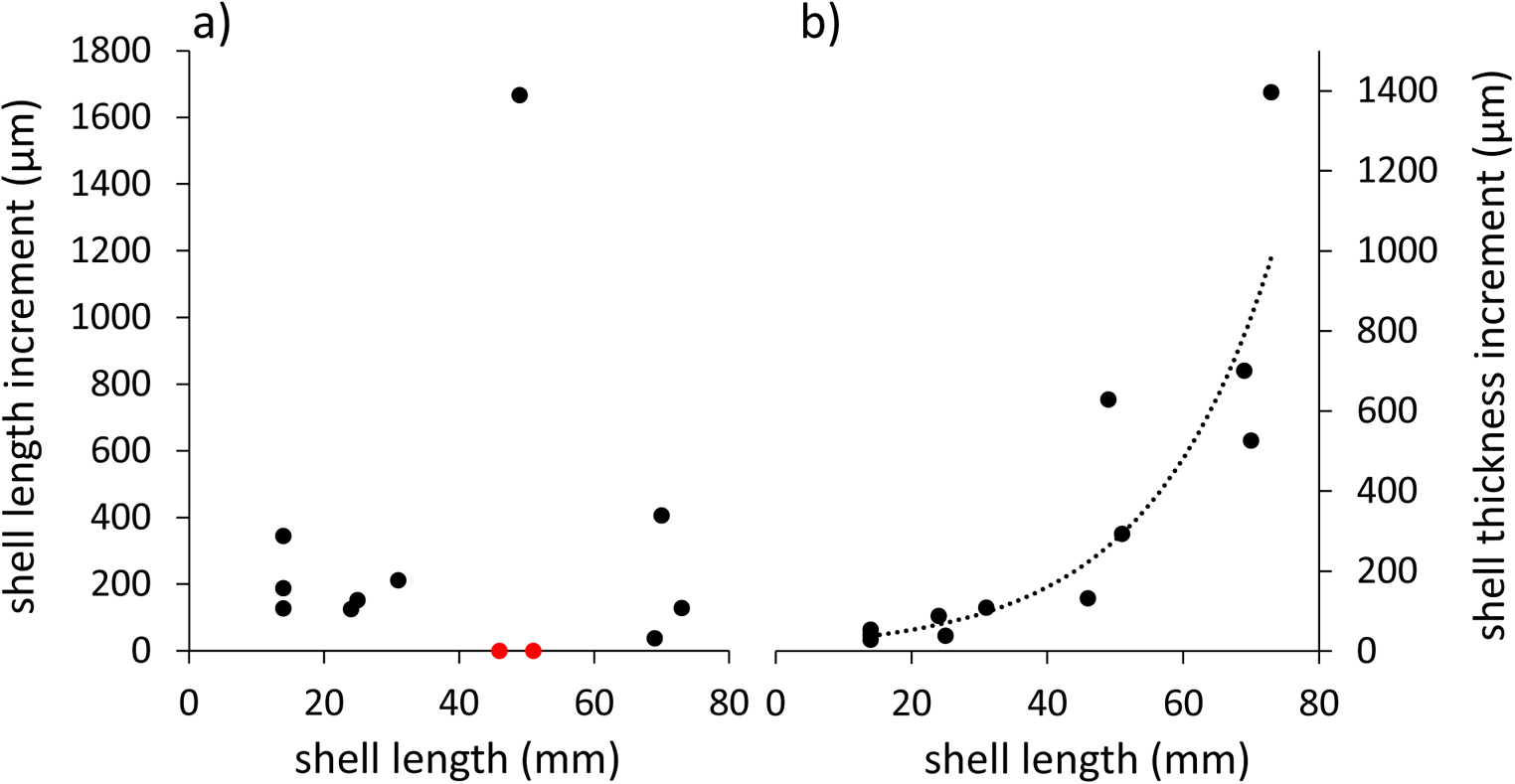
Batch 1 *G. childressi* shell length (SL, a) and shell thickness (ST, b) growth after 21 months, as deduced from calcein staining marks using a using a Leica M165 FC stereo microscope. Shell thickness increased exponentially increased with mussel size (b; ST increase (µm) = 17.737e^0.055x^, x = SL (mm), R² = 0.89).

#### Methane and oxygen consumption

Batch 1 mussels’ methane consumption rates ranged between 0.03 µmol g^−1^ h^−1^ and 0.17 µmol g^−1^ h^−1^, batch 1 oxygen consumption rates between 0.18 µmol g^−1^ h^−1^and 0.92 µmol g^−1^ h^−1^ (fig. 8a). There was a slight trend towards an increasing oxygen consumption of mussels at higher methane consumption rates (MO_2_ = 1.06(± 3.15, 95%CI) MCH_4_ + 0.33(± 0.28, 95%CI) [µmol g^−1^ h^−1^]; regression: R^2^ = 0.05, F = 0.56, p = 0.47; i. in fig. 8a). Methane consumption rates of batch 2 *G. childressi* that were cultured with methane supply (‘m’ and ‘a+m’) ranged between 0.1 µmol g^−1^ h^−1^ and 0.23 µmol g^−1^ h^−1^ in 2017 and had increased to a range between 0.17 µmol g^−1^ h^−1^ and 0.82 µmol g^−1^ h^−1^ in 2019 (fig. 8b). In 2017, oxygen consumption of batch 2 mussels increased significantly with higher methane consumption rates: MO_2_ = 2.19(± 1.78, 95%CI) MCH_4_ + 0.54(± 0.24, 95%CI) [µmol g^−1^ h^−1^] (regression: R^2^ = 0.55, F = 8.49, p = 0.023; ii. in fig. 8a).

**Figure 8.**
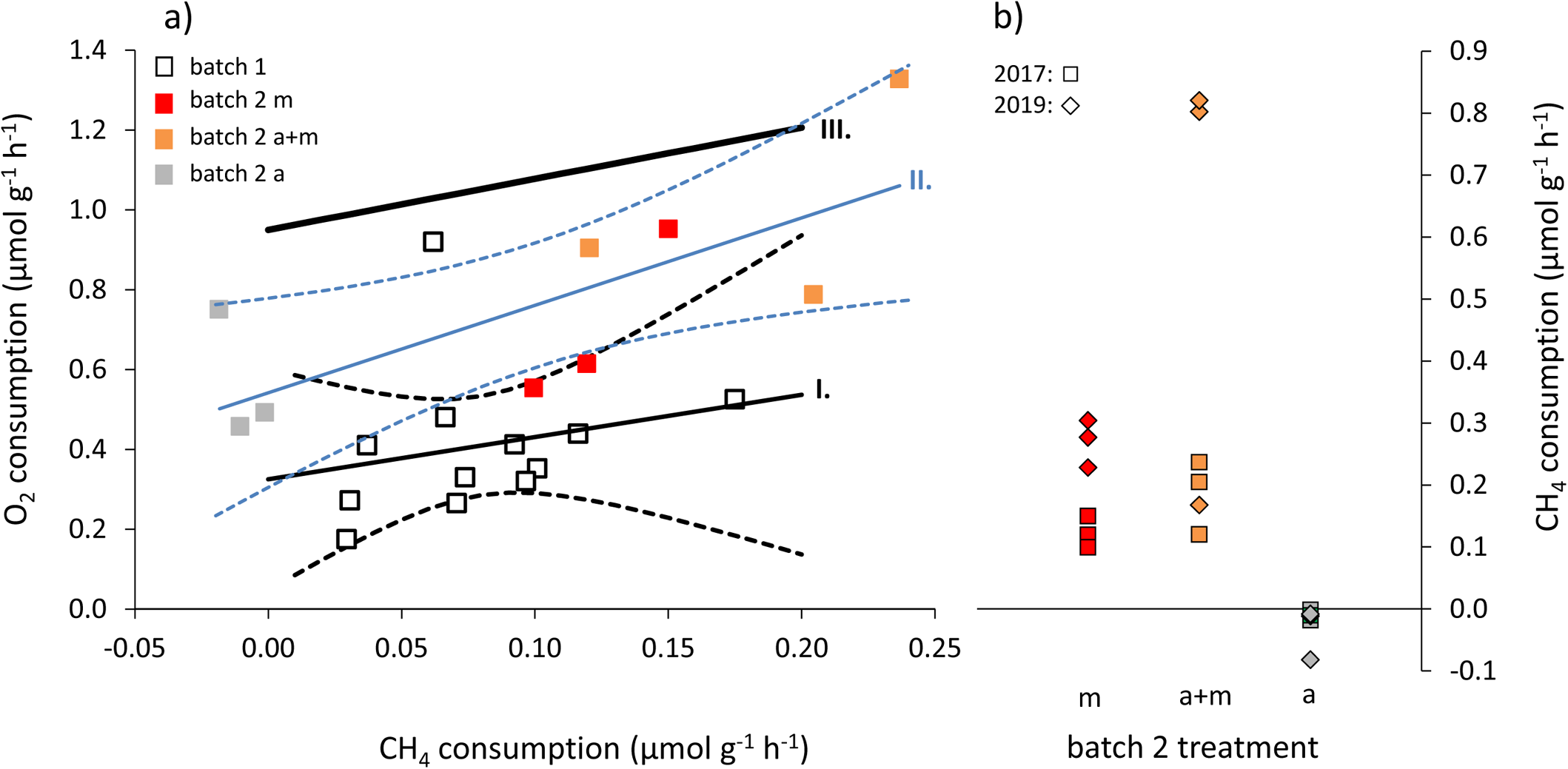
Gas consumption of batch 1 and 2 *G. childressi.* Oxygen consumption depending on methane consumption rate (a) with similar average slope but overall lower oxygen uptake of cultured batch 1 animals (I.) as compared to the relationship found by Kochevar et al. (1992, III.: MO_2_ (µmol g^−1^ h^−1^) = 1.19(±0.09, 95%CI) MCH_4_ (µmol g^−1^ h^−1^) + 0.95). Batch 2 mussels were cultured under 3 different nutrition regimes (CH_4_: ‘m’, algae + CH_4_: ‘a+m’, only algae: ‘a’; II.). Dashed lines in a) represent 95% confidence envelopes. Methane consumption of batch 2 mussels was measured twice, in 2017 and 2019 (b).

#### Activity and seawater methane concentrations

The number of daily valve gape hours of batch 2 mussels averaged across all three repeated runs of diurnal variations of seawater CH_4_ concentration (fig. 5, suppl. fig. 5) was independent of the nutritional treatments the *G. childressi* were previously acclimated to (fig. 9). However, as a tendency, deep-sea mussels that were acclimatized to CH_4_ reacted slightly differently to changes in seawater CH_4_ than mussels that were exclusively acclimated to microalgae food without CH_4_ addition. Overall, we found a trend towards longest activity phases of *G. childressi* at intermediate seawater CH_4_ concentrations (20-45 µM), and not at the highest concentration (57 µM, fig. 10). All mussels reduced valve gaping time periods in the absence of CH_4_ in seawater. However, this pattern - as well as the reduced activity at the highest CH_4_ level - was most prominent at *G. childressi* that were acclimated to CH_4_ (‘m’ and ‘m+a’, fig. 10a and 10b). The reactions of deep-sea mussels to changes in seawater CH_4_ appeared to be generally stronger when CH_4_ levels were gradually increased (suppl. fig. 6a-c) as to when the CH_4_ concentration was reduced (suppl. fig. 6d-f).

**Figure 9.**
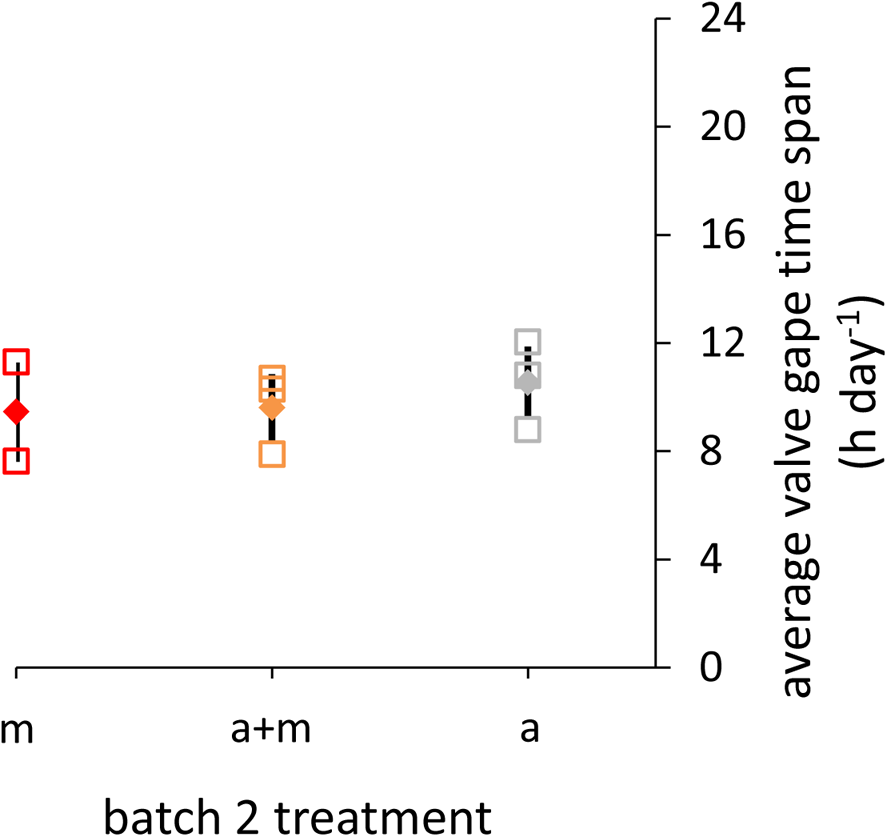
Average daily valve gape time of batch 2 *G. childressi* during the activity experiment based on time lapse image analysis. Mussels were cultured under 3 different nutritional regimes: CH_4_: ‘m’, algae + CH_4_: ‘a+m’, only algae: ‘a’. Diamonds: mean ± SD.

**Figure 10.**
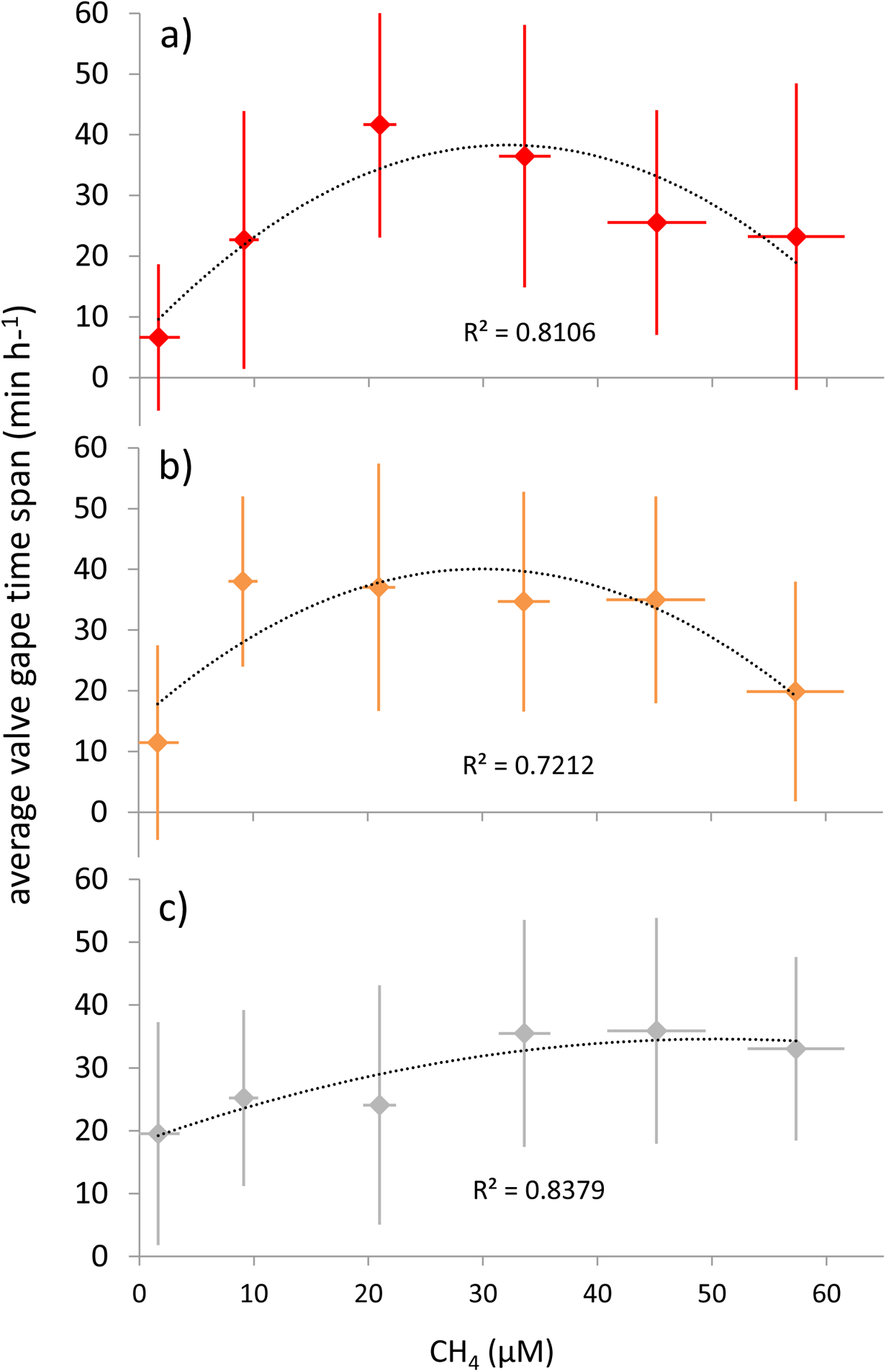
Valve gape activity of *G. childressi* mussels from different nutritional treatments (a: CH_4_ ‘m’, b: algae + CH_4_ ‘a+m’, c: algae ‘a’) in response to step-wise increasing and decreasing seawater CH_4_ concentrations (each 3 times) based on time lapse image analysis. Data are mean ± SD, n = 3.

#### Particulate food consumption

Batch 1 *G. childressi* cleared both, *Rhodomonas* sp. and *Nannochloropsis* sp. cells from seawater (fig. 11a), however, *Rhodomonas* sp. at a higher rate. Larger mussels (shell length 85 ± 7 mm) were characterized by higher phytoplankton clearance rates than smaller mussels (shell length 51 ± 3 mm). After being cultured for eight months under different nutritional scenarios, batch 2 *G. childressi* were characterized by different *Rhodomonas* sp. clearance behaviors: mussels acclimated to CH_4_ and microalgae food supply (‘a+m’) cleared *Rhodomonas* at highest rates of 91.1 mL h^−1^ (fig. 11b) which corresponds to 6.4 x 10^5^ cells h^−1^ (respectively 1.12 J h^−1^; at 1.75 µJ cell^−1^; Riisgård et al. 2012). *G. childressi* that were previously acclimated to CH_4_-free seawater (‘a’) cleared *Rhodomonas* sp. in the absence of CH_4_ - but not when the seawater was enriched with CH_4_ (fig. 11b).

**Figure 11.**
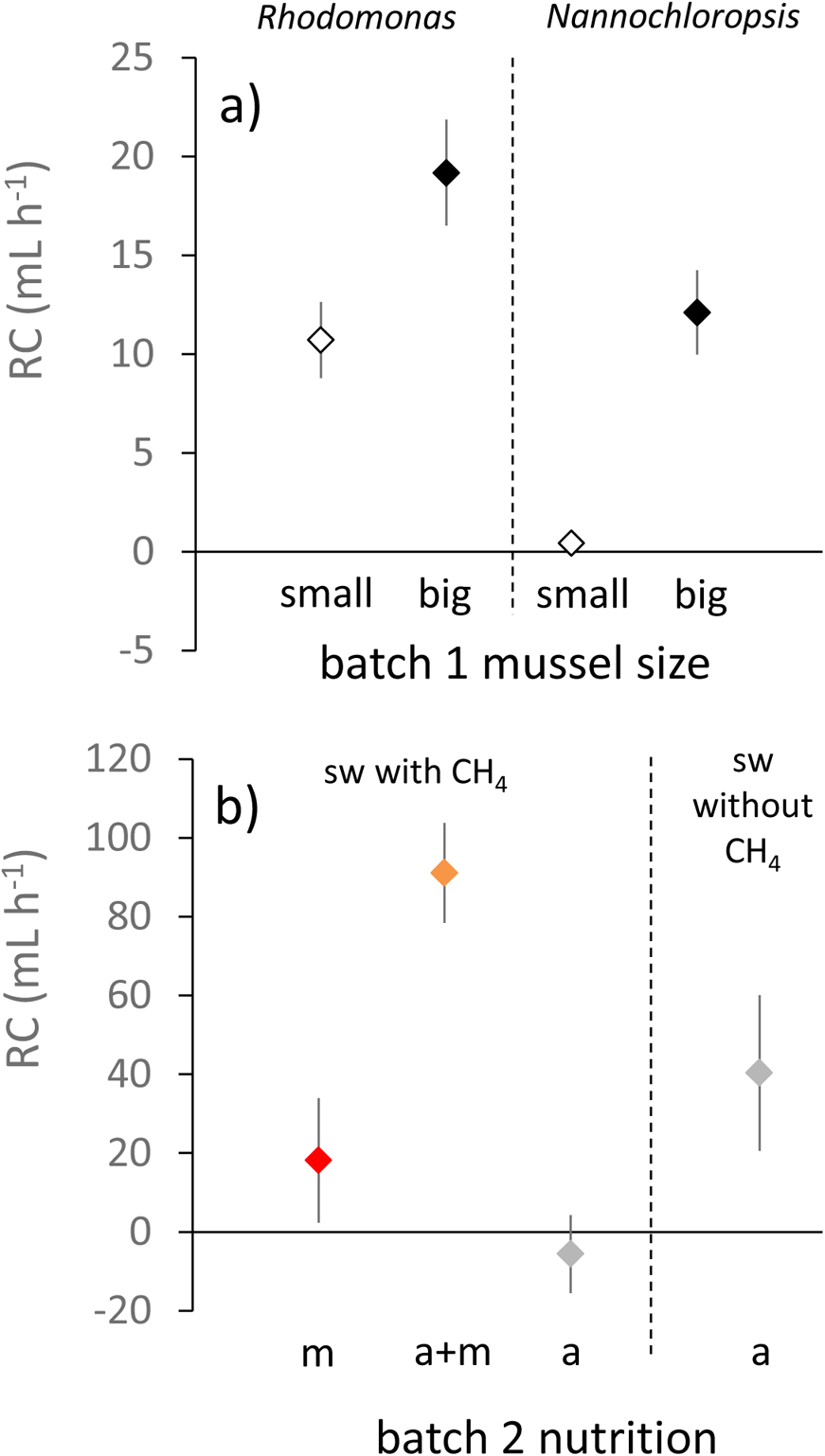
Clearance rates (RC) of *G. childressi* mussels filtering phytoplankton. (a) RC of different-sized batch 1 mussels (‘small’: 51 ± 3 mm SD, ‘large’: 85 ± 7 mm SD) that received either *Rhodomonas* or *Nannochloropsis* cells. (b) Clearance trials of batch 2 mussels from different nutritional treatments (CH_4_: ‘m’, algae + CH_4_: ‘a+m’, only algae: ‘a’) filtering *Rhodomonas* cells. The measurement of clearance rates of *G. childressi* cultured in the ‘a’ treatment (without CH_4_ supply) was conducted twice: once while mussels were placed in CH_4_-enriched seawater, and, subsequently, once without CH_4_. Data are mean ± SE.

#### Stable isotopes

Tissue stable isotope values of the analyzed *G. childressi* (δ^15^N: −2.57-1.22 and δ^13^C: −59.5- - 54.4) were well in the range of those of other *G. childressi* specimen collected in the Green Canyon area in the Gulf of Mexico (MacAvoy et al. 2008; Becker et al. 2010). However, all cultured mussels had a slightly reduced δ^13^C fractionation as compared to mussels sampled when they arrived in Kiel (without being influenced by our artificial nutritional treatments: ‘t_0_’, fig. 12b). This effect was again slightly stronger in *G. childressi* that were cultured with microalgae only (without CH_4_) when compared with mussels cultured with CH_4_ supply. This agrees with the lower δ^13^C fractionation of *Rhodomonas* sp. microalgae as compared to the CH_4_ used to equilibrate the aquaria. Annual carbon turnover rates, calculated as %Δδ^13^C_ann,_ were 7.9% for ‘a’-cultured *G. childressi* and 7.6% for ‘m’-cultured mussels. Finally, only *G. childressi* that were acclimated to a microalgae diet (‘a’) showed a weak tendency towards a positive δ^15^N fractionation (fig. 12a). The calculated annual N turnover rate (%Δδ^15^N_ann_) of ‘a’ cultured *G. childressi* was 3.9%.

**Figure 12.**
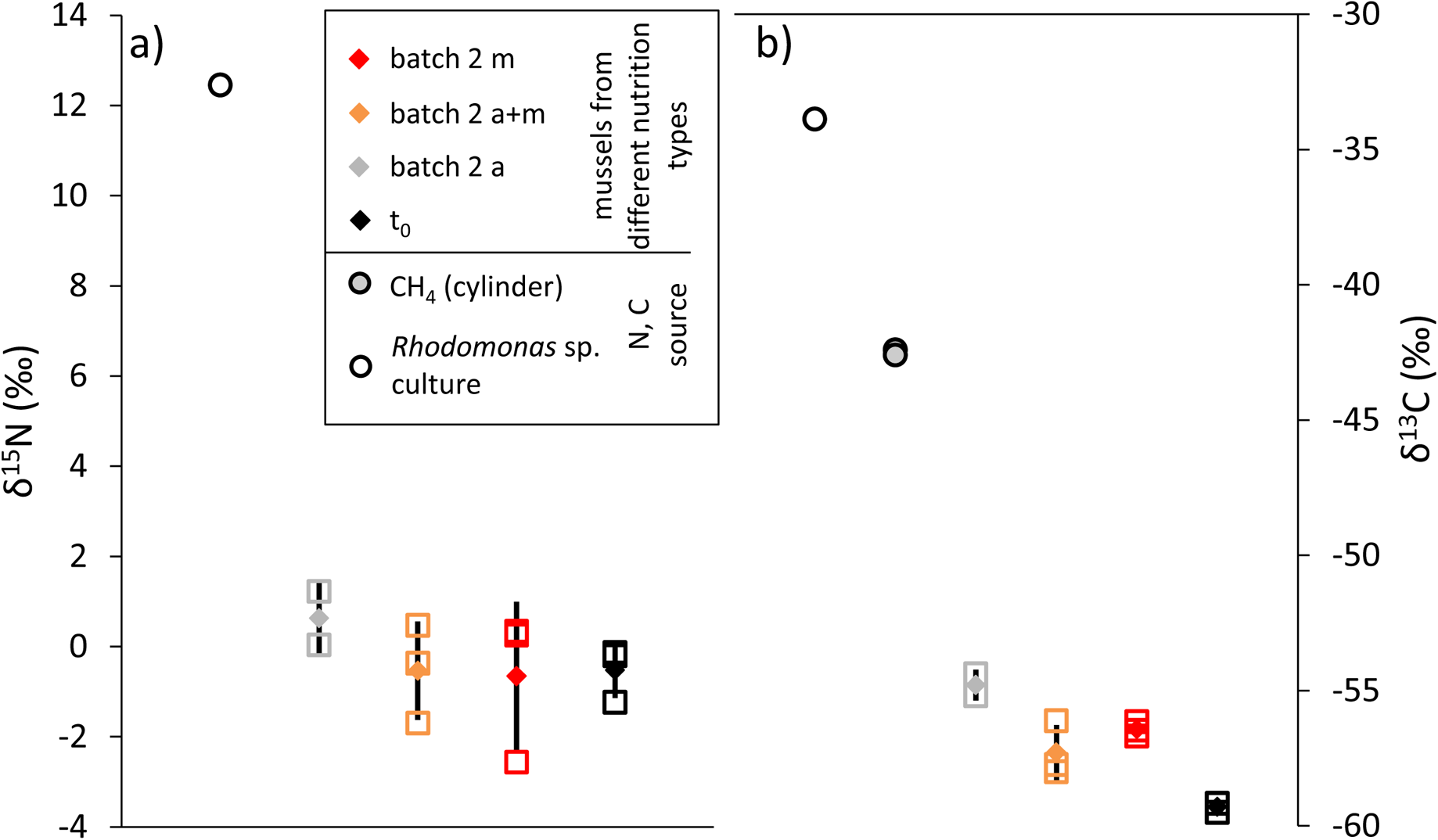
Stable isotopes (a: δ^15^N, b: δ^13^C) of batch 2 *G. childressi* that experienced different nutritional regimes (CH_4_: ‘m’, algae + CH_4_: ‘a+m’, only algae: ‘a’) in our culture. As reference, t_0_ animals were sampled upon arrival at GEOMAR after spending 2 months post-capture at Charles Fisher’s ‘deep-sea lab’ aquarium system, Pennsylvania State University. Diamonds: mean ± SD.

## 4. Discussion

Culture systems that allow for long-term studies of deep-sea organisms are difficult to realize and, hence, rare. Here, we present a recirculation aquarium system with an attached CH_4_-air gas mixing device, in which we can keep deep-sea mussels (*G. childressi*) for years in good condition. We also demonstrate that variation of seawater methane concentration impacts methane and oxygen consumption rates of experimental animals and show that acclimation of animals to different nutritional treatments influences whole-animal physiological properties. These pilot experiments indicate a strong potential for our culture system to enable exciting experimental work to better understand nutritional bio-energetics of symbiont bearing deep-sea mussels.

Overall, we were able to design and set-up a safe recirculating culture system in this proof-of-concept study to automatically supply deep-sea mussels with high methane levels for years. After overcoming technical limitations, namely sub-optimal production of the methane-air gas mixture by the gas mixing device in the beginning of the culture period (batch 1 animals), stable concentrations of 50-60 µM methane in seawater via equilibration with 4% methane in air gas mixtures could be maintained consistently (fig. 5, suppl. fig. 1). It is not possibly to produce higher methane concentrations using this system design, as the lower explosion limit for methane does not allow for equilibration with gas mixtures containing more than 4.4% methane.

During the time in our culture system, *G. childressi* acquired energy, both, via their methanotroph symbionts, which oxidize the methane provided into the culture system, and from microalgae added to the seawater. Both nutritional strategies could be corroborated by direct measurements of methane and microalgae uptake. The measured maximum methane (< 1 µmol g^−1^ h^−1^) and oxygen (< 1.5 µmol g^−1^ h^−1^) consumption rates of *G. childressi* cultured in our system were lower than reported by Kochevar et al. (1992; III. in fig. 8a), who, however, had a high variance in their data and measured at much higher seawater CH_4_ concentrations as well. Metabolic rates appeared to be strongly reduced in batch 1 mussels, as can be seen when comparing O_2_ consumption at similar CH_4_ consumption rates (offset of I. and III. in fig. 8a). Metabolic rates could be elevated by increasing CH_4_ supply about two-fold when culturing mussels of batch 2: O_2_ and CH_4_ consumption rates of mussels nourished with higher levels of CH_4_ for 4.5 months (II. in fig. 8a) were close to those measured by Kochevar et al. (1992). Furthermore, after another 2 years of culture, the batch 2 CH_4_ consumption rates had increased substantially (fig. 8b), especially of those mussels that also received microalgae food (‘a+m’). Lower methane consumption rates are likely due to a partial loss of methanotroph endosymbionts in the gills during transfer from the deep sea and culture under insufficient methane supply (Kádár et al. 2005). Batch 1 and 2 animals arrived in our lab 0.5 and 2 months post-capture, respectively, and we presume that methane supply in cultures post-capture and pre-arrival to Kiel (unquantified) was sub-optimal. Previous *in situ* work demonstrated that bathymodiolid mussels can rapidly lose most endosymbionts within weeks (Détrée et al. 2019). Thus, future culturing efforts should utilize experimental animals rapidly transferred from the site of capture to our culturing system to prevent this initial loss in symbiont density and diversity.

The observed up to 4-fold increase in methane uptake rate of batch 2 mussels may be explained by a subsequent re-acquisition of methanotrophs under our improved culture system conditions (Kádár et al. 2005; Riou et al. 2008; Tietjen et al., unpublished results). Deep-sea mussels that were fed only with microalgae apparently lost all methanotroph symbionts, as indicated by a lack of methane consumption after an acclimation time of 3.5 months to this regime (Fig. 8b). While microalgae-fed animals could maintain a relatively high condition, they were characterized by a much lower survival than methane acclimated groups, indicating the ultimately unsustainability nature of this nutritional treatment. These mussels’ valve gape activity was relatively unaffected by methane concentration during our behavior monitoring trials (fig. 10). However, they filtered *Rhodomonas* sp. from the seawater only in the absence of methane, indicating a fundamental change in whole organism physiology and behavior elicited by our acclimation treatment. This was a surprising observation as the ventilation rate of *G. childressi* – unlike that of *M. edulis* – is assumed to be independent of stimuli by the filtered particles in suspension (Page et al 1990).

Also, to our knowledge, no direct methane-sensing organ has been described for deep-sea mussels so far. However, valve gape activity of mussels cannot be directly translated into ventilation or filtration rate, as gill-cilia driven water convection rate can be controlled independently of valve gape (Riisgård et al. 2011). On average, *G. childressi* valve gape opening probability for CH_4_-supplied experimental groups increased when methane concentration was gradually increased with a maximum at intermediate values (20-45 µM). We assume that this preference of intermediate methane concentrations can be attributed to an increase in metabolic activity of the methanotroph endosymbiont bacteria with rising methane concentrations and a saturation of their methane requirements at high seawater methane concentrations. How methane demand of endosymbionts and water convection rate by the mussel are coordinated and regulated, is unclear and can be explored in future experiments using our culturing system.

Besides a reasonably good condition index of *G. childressi* cultured in our recirculation aquarium system, almost no shell length growth and only low shell thickness growth could be detected. In comparison, shell length increments of several mm per year have been observed in the field at a seep nearby the sampling station our animals originated from (GC 234; Nix et al. 1995). The methane concentrations in our cultivation system - which are still lower than measured at many cold seep sites - are unlikely to be the main reason for the lower growth rates: Nix et al. (1995) measured similar or even lower methane concentrations at sites where much higher shell length growth rates of *G. childressi* were observed. Furthermore, *G. childressi* in our culture were in good body condition, able to invest surplus energy into shell thickness growth and, hence, not starving. However, our measured methane consumption rates suggest that the mussels lost a number of their methanotroph symbionts during transfer from the deep sea to our culture system – and that these were only partially able to re-grow. A higher number of ‘farmed’ symbionts would certainly allow experimental animals to invest more energy into shell (length) growth.

However, whole soft tissue stable isotope data clearly indicates that both nutritional strategies, (consumption of farmed endosymbionts, consumption of microalgae cells) were utilized by our cultured mussels. While the δ^13^C of −59.3‰ of mussels sampled upon arrival at GEOMAR (‘t_0_’) indicates a significant input of biogenic methane (typically < −60‰) at the sampling site GC249, the lower C isotope fractionation of the cultured *G. childressi* mussels (m_c_: ‘m’, ‘a+m’ and ‘a’) can only be explained by uptake of thermogenic methane (from gas cylinders: δ^13^C = −42.5‰) and / or microalgae (*Rhodomonas* sp.: δ^13^C = −34‰; Brooks et al. 1987; Kennicutt et al. 1992; Becker et al. 2010). The obvious delta of mussel soft tissue δ^13^C and the δ^13^C of the C sources (thermogenic methane, respectively *Rhodomona*s sp.), Δδ^13^C, though, can be interpreted as incomplete soft tissue turnover (Dattagupta et al. 2004). Annual C turnover (<10% for ‘a’ and ‘m’ nutritional groups) are significantly lower than the 40% – 78% Dattagupta et al. (2004) calculated for G. *childressi* that were transplanted in the field and displayed strong shell growth – and also than the 35% – 53% the authors calculated for transplanted mussels that did not grow after transplantation. Thus, metabolism and growth of *G. childressi* living in our deep-sea recirculation aquarium system appears to have been reduced.

While post-capture loss and reduced density of symbionts during most of the culture time within our culture system may partially explain the low growth rates observed, low hydrostatic pressure might be another contributing factor. (Severe) depressurization damage to DNA and shell integrity has been described, however, predominantly for bathymodiolids collected from depths >1000 m (Dixon et al. 2004, Kadar et al. 2008). Dixon et al. (2004) suggest a physiological barrier for populations from depths >1000-1500 m that prevents successful acclimation to surface pressures. For *G. childressi* that live in shallower habitats (<1000 m), Arellano & Young (2009, 2011) demonstrated that gamete release and development of viable larvae can be induced at surface pressure. These findings, in combination with our results, indicate that culturing efforts under surface pressure may be sufficient to study a range of important biological processes in *G. childressi* from relatively shallow deep-sea habitats. Studies utilizing shallow water invertebrates could demonstrate significant physiological plasticity during acclimation to pressures resembling water depths of ca. 1000 m, with changes in membrane composition (‘homeoviscous adaptation’) likely contributing to survival at elevated pressures (Brown et al. 2019). Thus, plastic re-organization of cellular structures might enable some marine invertebrates to acclimate to pressures in the 1-100 bar range in both directions. However, comparative growth trials of experimental *G. childressi* at ambient vs. high pressure (50-80 bar) are needed to quantify the impacts of hydrostatic pressure variation on fitness. Thus, a further development of the described culture system to support long-term experiments under pressures of up to 80 bar would be needed. Further, meaningful ecological studies will need multiple culturing systems with separate filtering and methane sensing systems for replicated measurements and factorial experiments. Due to the relatively low costs of methane sensing equipment, this seems a realistic option for the near future (Bastviken et al. 2020).

### Conclusions & outlook

We have shown that our newly designed recirculating culture system is suitable for long-term maintenance experiments with cold seep colonizing *G. childressi,* at seawater methane concentrations that can be found in natural deep-sea habitats of this species. Nutritional regimes involving controlled microalgae and continuous methane addition enabled good maintenance of body condition and some shell growth. Long-term acclimation of *G. childressi* to relatively high methane concentrations (ca. 50-60 µM) can increase methane uptake rates of experimental animals, indicating that symbiont depleted animals may be able to increase symbiont density again. On the other hand, we show that acclimation to a microalgae diet without addition of methane leads to phenotypes that do not consume methane anymore, suggesting loss of symbionts.

Thus, with the caveats mentioned above in mind (potential depressurization pathologies), a number of exciting questions can be addressed with the described system if it is properly replicated. First, experimental modulation of symbiont density through seawater methane supply will enable fundamental studies related to nutritional contributions of particulate food vs. energy and nutrients derived from consumption of symbionts to the host mussel energy budget.

Metabolic exchange processes between symbiotic partners can be studied, as well as methane sensing processes of mussel host and symbiont and how they drive water convection activity of the host to supply methane to symbionts. Arellano & Young (2009) demonstrated successful generation of *G. childressi* larvae in the laboratory, but were only able to rear larval stages to the first shell forming stage. Future studies can build on this work and try to maintain mussels in good condition for reproduction, and then spawn parental animals to raise larval stages ideally to settlement. Such data is urgently needed to better constrain biophysical models to estimate connectivity of deep-sea mussel populations (Breusing et al. 2016). Key parameters, such as duration of larval development, swimming speed and behavior, as well as chemotactic abilities are unknown at present. Methane sensing abilities of larvae and adults could be studied in detail using our culturing system by creating methane gradients in behavioral trials. Currently, nothing is known about methane sensing abilities of *G. childressi*, but it is likely that unknown sense organs exist to guide larvae to seep sites once they are competent for settlement and adults to locations with optimum methane supply. Thus, we hope that culture systems as described here will stimulate more research to in detail investigate how these fascinating mussel holobionts function and how they can be protected in a deep-sea environment that is increasingly exploited.

## 5. Acknowledgements

We thank Nicole Dublier and Chuck Fisher for collection and transport to Germany of batch 1 mussels and Iliana Baums, Samuel Vohsen and, again, Chuck Fisher for collecting batch 2 mussels, giving them a temporary home at the Chuck Fisher deep-sea lab and sending them to GEOMAR and Ulrike Panknin for culturing *Rhodomonas*. This study was supported by the Cluster of Excellence ‘The Future Ocean’ project ‘Kiel Marine Organism Culture Centre – KIMOCC’, funded by the German Research Foundation (Deutsche Forschungsgemeinschaft) DFG.

