## supplementary tables and figures for "Long-term culture system for deep-sea mussels *Gigantidas childressi*"

**Supplementary table 1.** Overview on Bathymodiolidae cultures in other laboratories.

| species | laboratory | methane supply | sulphide supply | other nutrition | culture duration according to source | source |
|---|---|---|---|---|---|---|
| *Bathymodiolus azoricus* | LabHorta, Azores | Continuous $CH_4$ bubbling of aquariums, creating approx. 70 µM | 20 mM sulphide solution dispensed for 15 min every hour, creating sulphide concentra-tions of 1-90 µM | Algae paste of a mixture of *Nannochloropsis* sp. and *Tetraselnis* sp. | > 12 months | Colaço et al. 2006 |
| | | Continuous $CH_4$ bubbling of aquariums, creating approx. 8-92 µM | 20 mM sulphide solution dispensed for 15 min every hour, creating sulphide concentra-tions of 1-30 µM | - | ? | Riou et al. 2008 |
| | | - | - | freeze-dried ocean plankton enriched with vitamins (Ocean Plankton, Hikari BIO-PURE FD) | 3 months | Bettencourt et al. 2011 |
| ‘B*athymodiolus*‘ = *Gigantidas childressi* | Oregon Institute of Marine Biology | Diurnal placing of mussels into ‚feeding‘ tank with $CH_4$-bubbled seawater for 3 hours | - | - | 6 months | Arellano & Young (2009) |
| | ‘C. Fisher deep-sea lab‘, Pennsylvania State University | $CH_4$-saturated seawater led to the bottom of mussel aquarium | - | - | several (?) years | pers. comm. Samuel Vohsen |

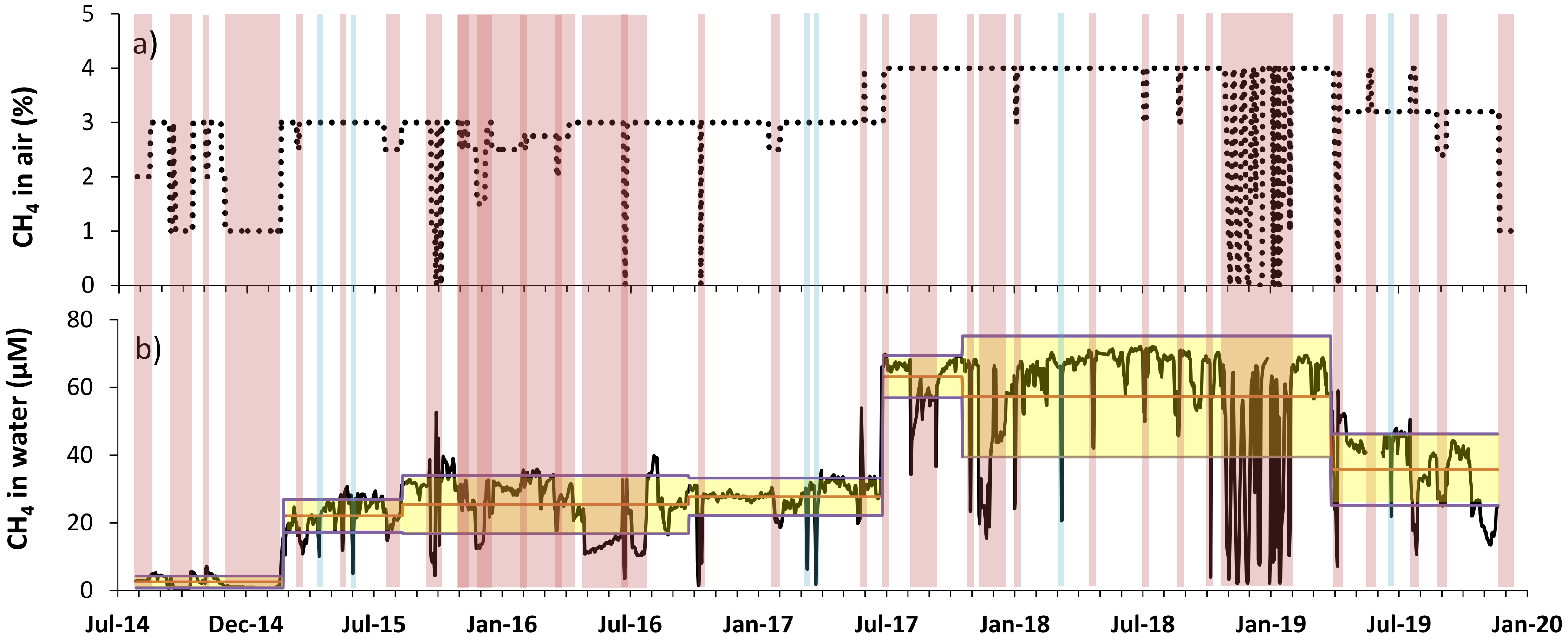


**Supplementary figure 1.** Methane supply to the deep-sea mussel culture system over time (July 2014 – Jan 2020). A): Proportion of $CH_4$ in air bubbled into the culture seawater, b): $CH_4$ concentration in seawater with average concentrations of different time periods (yellow area: SD during the respectively time period). Pink bars mark variations of $CH_4$ concentration caused by intended changes of the system, namely changes in %$CH_4$ in air, changes in gas flow or other experiments. Blue bars mark variations of $CH_4$ concentration caused by technical malfunctions: air-gas mixture leakages, gas supply problems or power failures.

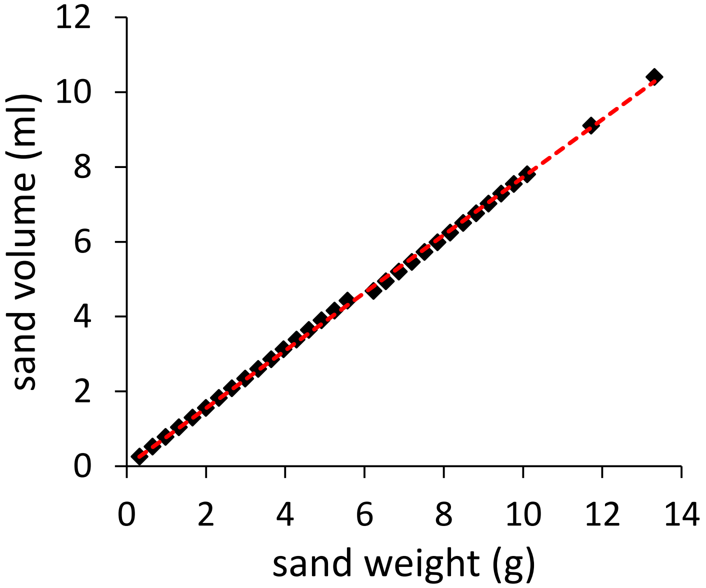


**Supplementary figure 2**. Volume (V) – weight (m) relationship of silica sand (grading 50-1000 µm) used to measure the mussel shell volume. A standardized cup of exact 0.2602 mL was used to measure 1 to 30 as well as 35 and 40 standardized units of sand. Equation: V [mL] = 0.7728 [g mL$^{-1}$] m [g], $R^2$ = 0.9993, n = 32.

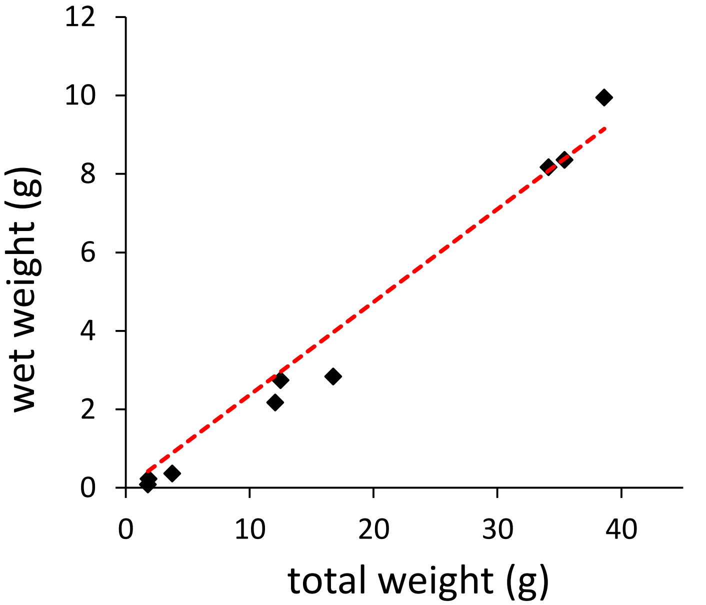


**Supplementary figure 3**. Mussels wet weight – total weight relationship of *G. childressi* for $C_i$ measurement. Equation: wet weight [g] = 0.2369 x total weight [g], $R^2$ = 0.9763, n = 9.

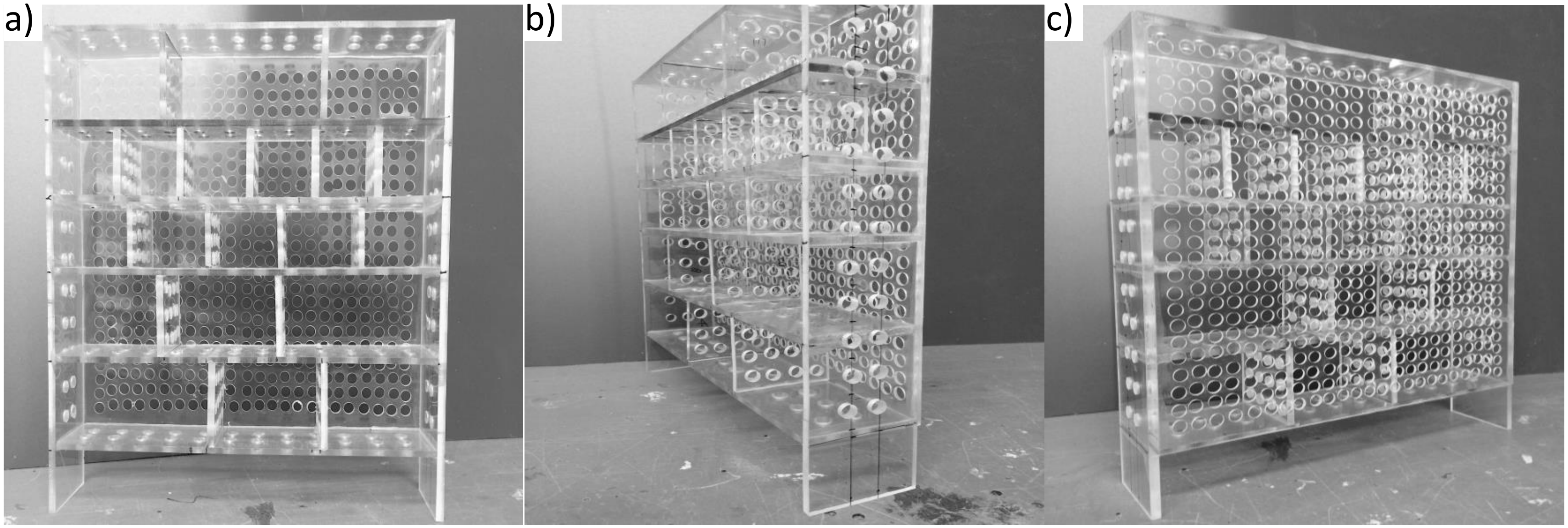


**Supplementary figure 4**. Acrylic glass rack used to observe *G. childressi* reactions to changes in seawater methane concentrations. After mussels were set individually in a compartment the rack was placed int the aquarium with the front side (a) facing tightly to the aquarium wall. B) and c) show holes drilled into rack's walls to allow freshly aerated seawater to reach the mussels. Mussels could be observed with an IR-camera while seawater methane concentrations were lowered or raised.

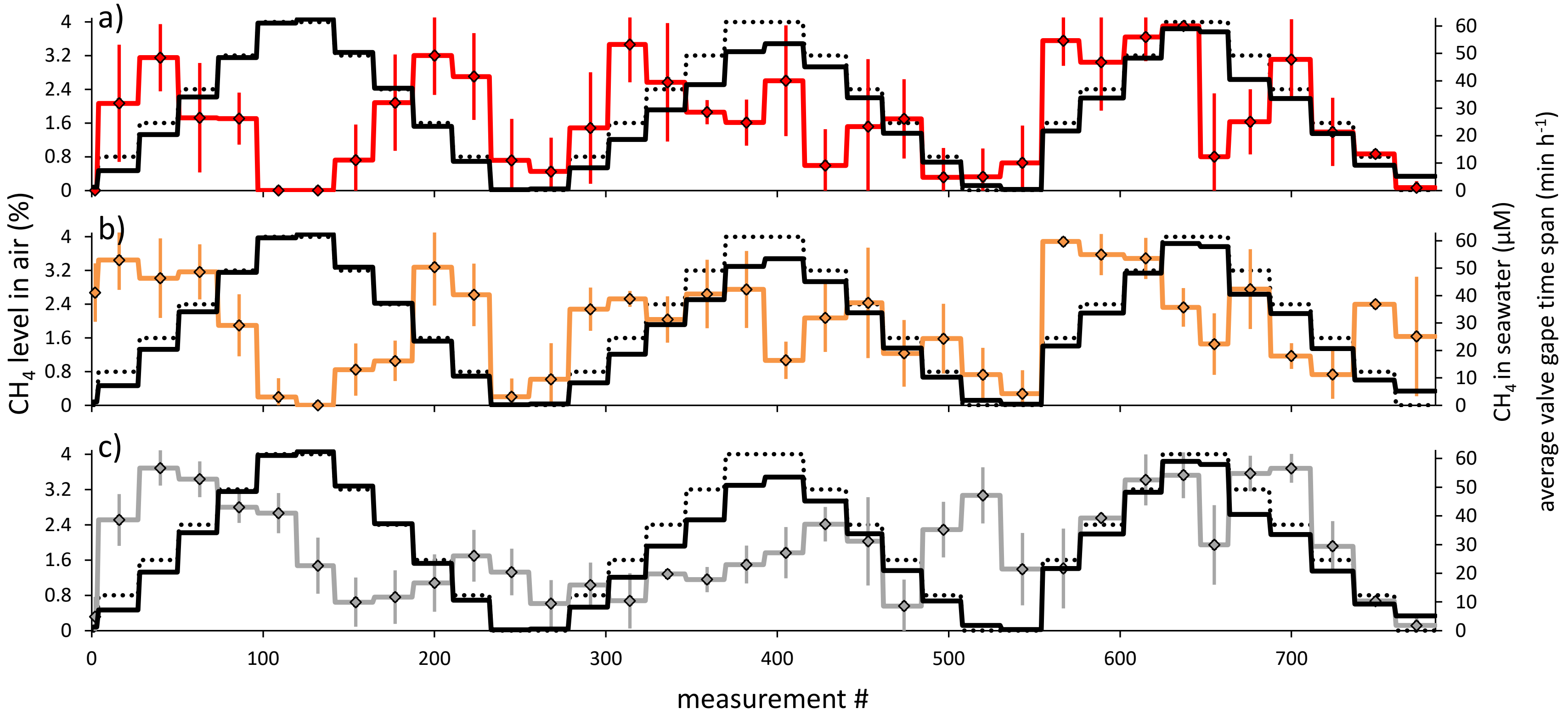


**Supplementary figure 5**. Valve gape activity of *G. childressi* from different nutrition treatments (a: $CH_4$, b: algae + $CH_4$, c: algae) at different stepwise changed $CH_4$ concentrations. Black solid line: seawater $CH_4$ concentration, black dotted line: $CH_4$ level in bubbling air. Data are mean ± SD.

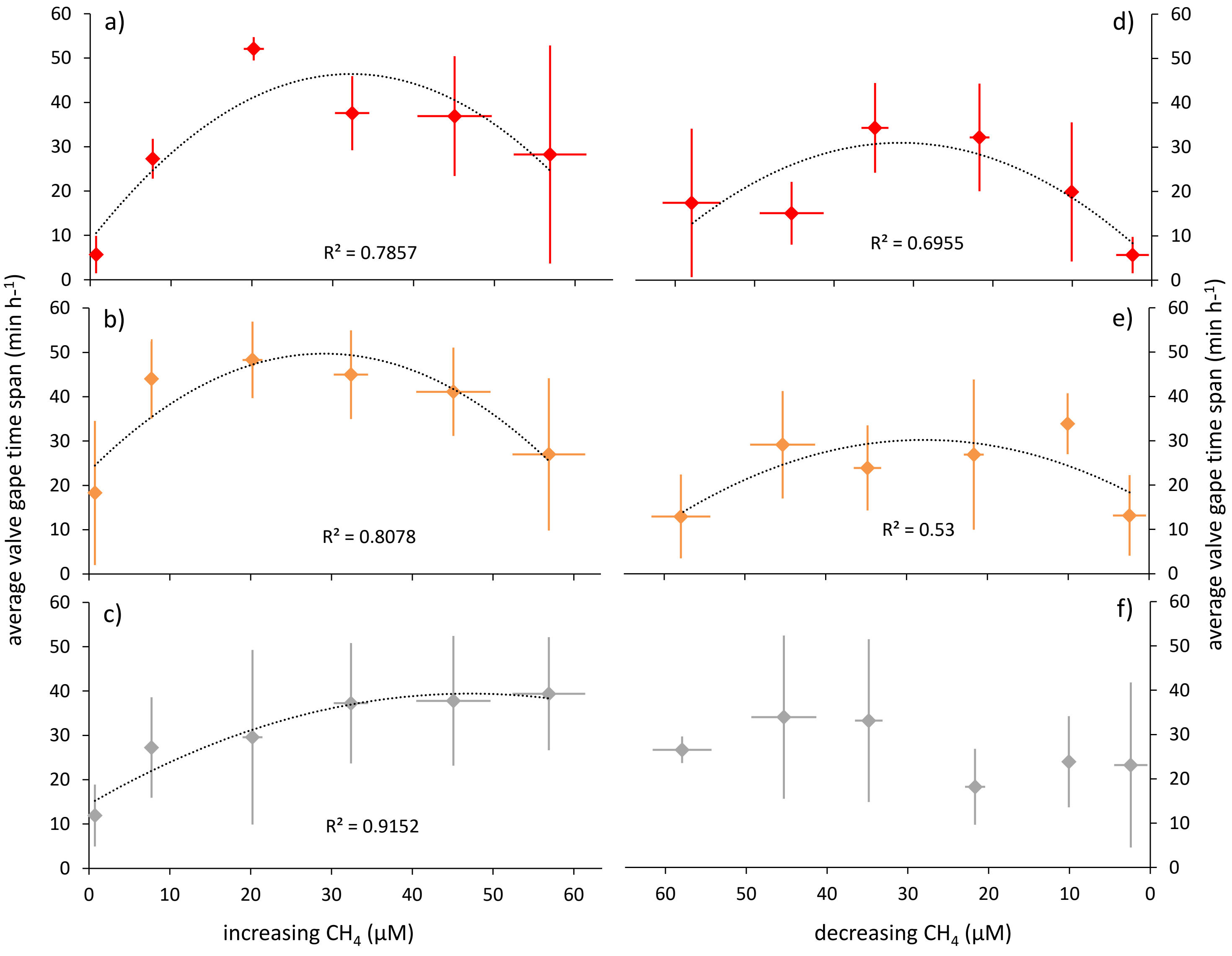


**Supplementary figure 6**. Valve gape activity of *G. childressi* mussels from different nutrition treatments (a, d: $CH_4$, b, e: algae + $CH_4$, c, f: algae) at increasing (a-c) or decreasing (d-f) seawater $CH_4$ concentrations. Data are mean ± SD.

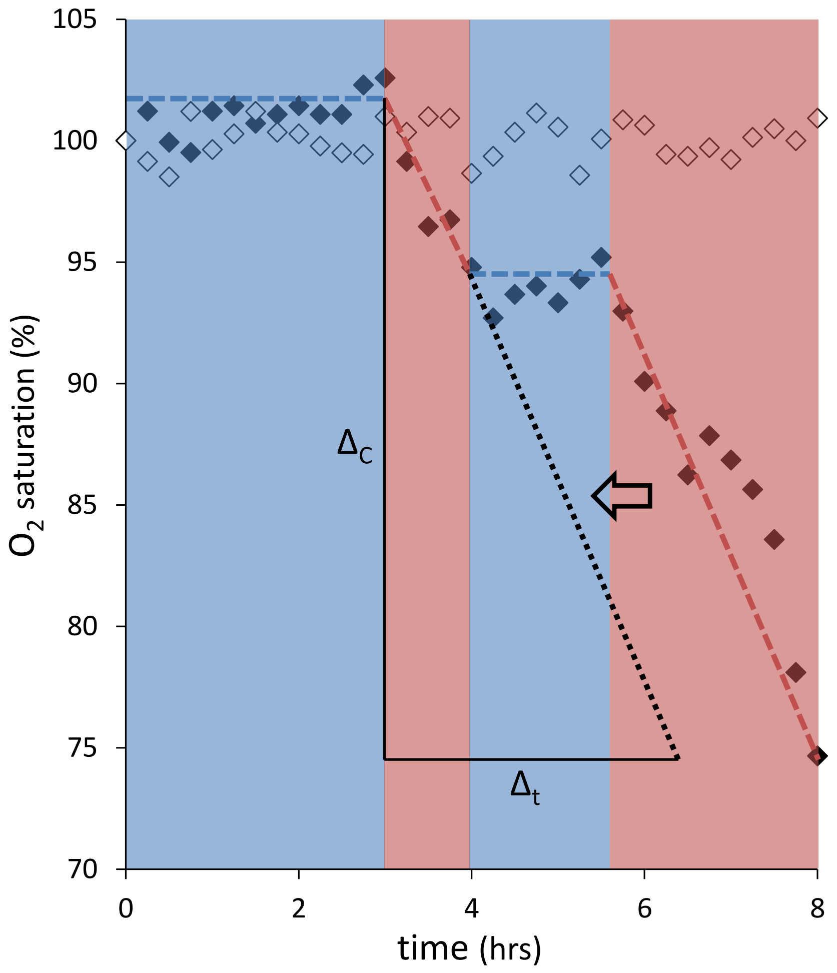


**Supplementary figure 7**. Assessment of *G. childressi* respiration time ($\Delta_t$) during oxygen measurements. Periods of stable (or increasing) $O_2$ saturation (blue areas) were addressed as times of mussel shell closure and cessation of gas exchange and, hence, ignored in the assessment of $\Delta_t$. Frequency of measurements: 4 $h^{-1}$.
